# Causal-inference regulates audiovisual spatial recalibration via its influence on audiovisual perception

**DOI:** 10.1101/2021.03.16.435619

**Authors:** Fangfang Hong, Stephanie Badde, Michael S. Landy

## Abstract

To obtain a coherent perception of the world, our senses need to be in alignment. When we encounter misaligned cues from two sensory modalities, the brain must infer which cue is faulty and recalibrate the corresponding sense. We examined whether and how the brain uses cue reliability to identify the miscalibrated sense by measuring the audiovisual ventriloquism aftereffect for stimuli of varying reliability. Visual spatial reliability was smaller, comparable to and greater than that of auditory stimuli. To adjust for modality-specific biases, visual stimulus locations were chosen based on perceived alignment with auditory stimulus locations for each participant. During audiovisual recalibration, participants were presented with bimodal stimuli with a fixed perceptual spatial discrepancy; they localized one modality, cued after stimulus presentation. Unimodal auditory and visual localization was measured before and after the recalibration phase. We compared participants’ behavior to the predictions of three models of recalibration: (a) Reliability-based: each modality is recalibrated based on its relative reliability—less reliable cues are recalibrated more; (b) Fixed-ratio: the degree of recalibration for each modality is fixed; (c) Causal-inference: recalibration is directly determined by the discrepancy between a cue and its final estimate, which in turn depends on the reliability of both cues, and inference about how likely the two cues derive from a common source. Vision was hardly recalibrated by audition. Auditory recalibration by vision changed idiosyncratically as visual reliability decreased: the extent of auditory recalibration either decreased monotonically, first increased and then decreased, or increased monotonically. The latter two patterns cannot be explained by either the reliability-based or fixed-ratio models. Only the causal-inference model of recalibration captures the idiosyncratic influences of cue reliability on recalibration. We conclude that cue reliability, causal inference, and modality-specific biases guide cross-modal recalibration indirectly by determining the perception of audiovisual stimuli.

**Author summary:** Audiovisual recalibration of spatial perception occurs when we receive audiovisual stimuli with a systematic spatial discrepancy. The brain must determine to which extent both modalities should be recalibrated. In this study, we scrutinized the mechanisms the brain employs to do so. To this aim, we conducted a classical recalibration task in which participants were adapted to spatially discrepant audiovisual stimuli. The visual component of the bimodal stimulus was either less, equally, or more reliable than the auditory component. We measured the amount of recalibration by computing the difference between participants’ unimodal localization responses before and after the recalibration task. Across participants, the influence of visual reliability on auditory recalibration varied fundamentally. We compared three models of recalibration. Only a causal-inference model of recalibration captured the diverse influences of cue reliability on recalibration found in our study, and this model is able to replicate contradictory results found in previous studies. In this model, recalibration depends on the discrepancy between a cue and its final estimate. Cue reliability, perceptual biases, and the degree to which participants infer that the two cues come from a common source govern audiovisual perception and therefore audiovisual recalibration.

## Introduction

### Cross-modal integration

In our daily lives, whenever we make a perceptual decision, we have to estimate properties of the environment such as the location of a barking dog. Usually multiple sensory cues for each property arrive in the brain. For example, a glimpse of the dog’s wagging tail and the barking both give away its location. However, due to external noise in the environment and internal noise in our sensory systems, the cues hardly ever lead to the very same percept, which renders the integration of cues from multiple sources challenging. How do we form a coherent percept despite these unavoidable cue conflicts? We do so by relying on a weighted mixture of the cues. This strategy becomes evident when not only the internal cues but also the external signals are in conflict. For example, when a ventriloquist speaks without moving his or her lips, there are two conflicting signals about the spatial location of the sound source. The auditory signal indicates that the sound originates from the ventriloquist, while the visual signal indicates that the sound comes from the “dummy”. In this case, vision dominates the combined spatial estimate of the sound source. This phenomenon is called the ventriloquism effect, a commonly used example to demonstrate “visual capture” in spatial localization [1–3]. However, when visual reliability, the precision with which the cues arriving in the brain match the external signal, is degraded enough to be lower than auditory reliability, the combined spatial estimate is no longer dominated by the visual but instead by the auditory signal, indicating that cue integration depends on the relative reliability of the cues [4–6]. This integration strategy is called reliability-based cue combination and it is computationally optimal in that it maximizes the precision of the resulting estimate. In addition to audiovisual spatial perception, reliability-based cue integration has also been found when integrating visual and auditory information to determine temporal rate [7–9], visual and haptic information to determine an object’s size and shape [10–12] and the perception of sequences of events [13], and visual and vestibular information to determine heading direction and to perceive own-body rotation [14–19].

However, it is not always the case that signals from multiple modalities are integrated. To continue with the ventriloquism example, if the speech sounds originate from someone else standing behind the stage or if the dummy’s mouth moves asynchronously relative to the speech sounds, the audience is unlikely to experience a ventriloquism effect. In other words, audiovisual integration breaks down when the two cues are too far apart either spatially or temporally to be perceived as coming from a common source. This breakdown of integration with spatial and temporal cue conflicts has been found not only in auditory-visual integration [20–25], but also for integration of visual and vestibular heading-direction signals [26, 27] and integration for visual and haptic surface thickness [28]. These findings suggest that the brain infers the causal relationship underlying cues from multiple modalities to derive the final percept.

Körding and colleagues (2007) formulated a model that describes how a Bayesian observer would determine the final percept using causal inference. The observer computes estimates of the underlying property given both causal scenarios, a common cause and different causes. The integrated estimate, for the scenario of a common cause, is based on the relative reliability of the sensory cues and a priori knowledge about the frequency of stimulus locations in the environment. The segregated estimate, for the scenario of different causes, is based on the relative reliability of each sensory cue and a priori knowledge. The observer further derives the probability of the two cues coming from a common cause and takes this probability into account when computing the final estimate from the two intermediate estimates, the integrated one and the segregated one. The causal-inference model accurately captures the nonlinear integration of audiovisual spatial signals by humans [29] and has been widely used to explain cross-modal perception in various contexts [26, 27, 30–39].

### Cross-modal recalibration

Reliability-weighted sensory cue integration maximizes the precision of the final estimate but not its accuracy; estimates are dominated by the more precise not by the more accurate cue. This preference of precision over accuracy might be rooted in the fact that single sensory signals can contain information about their reliability [40], but information about the accuracy of the sensory signals is impossible to derive from the signals themselves. Only systematic disparities between two sensory cues indicate a problem with the accuracy of the cues and, at the same time, provide a chance to address this issue. In this case, the brain should determine that either one or both senses are miscalibrated. It should adapt the interpretation of the cues to reduce that disparity and so, hopefully, maximize accuracy as well as precision. To continue with the ventriloquism example, after repeated exposure to spatially discrepant audiovisual stimuli, the shift in auditory spatial perception toward the visual stimulus still influences subsequent unisensory auditory localization when the visual signal is no longer available. This persisting shift is called the ventriloquism aftereffect, and has been replicated in many studies and across different modalities [31, 41–50]. In those ventriloquism studies, vision serves as the “teaching signal”, which is used to calibrate auditory or tactile perception when there is a consistent spatial cue conflict. This is a reasonable assumption given that visual information is usually more accurate and reliable than auditory information in terms of localizing objects spatially [51]. It remains unknown whether by default vision serves as the “teaching signal” in the spatial domain regardless of its reliability or whether visual perception can also undergo recalibration when its spatial reliability is greatly reduced. Only a few studies have tested the recalibration of visual space by audition. Some studies found systematic shifts in visual localization towards the direction of the auditory stimulus after repeated exposure to spatially discrepant audiovisual stimulus pairs, but the aftereffects were not as robust as the recalibration of auditory spatial perception by discrepant visual information [46, 52, 53]. Thus, vision might not always serve as the “teaching signal” and might undergo recalibration, but the evidence is weak.

### Reliability-based cross-modal recalibration

Which modality serves as the “teaching signal” in cross-modal recalibration might be determined based on cue reliability. More robust evidence for recalibration of visual perception was found in studies that examined the aftereffects of spatial visual-haptic and visual-proprioceptive as well as temporal visual-auditory cue discrepancies [54–59]. All of these studies found recalibration of visual perception when testing environmental properties for which the visual signal was not as reliable as the other signal, which suggests that the brain can use cue reliability when identifying miscalibrated senses. Burge and colleagues directly tested the relationship between cue reliability and the amount of recalibration by manipulating the reliability of the visual stimulus for slant to be either smaller, almost equated, or greater than that of the haptic cue [60]. As the visual cue became less reliable, greater recalibration of vision and less recalibration of haptics was observed, again supporting the notion that stimulus reliability plays a role in the process of visual-haptic recalibration just as it does in integration. The authors concluded, in agreement with others [53, 58], that each sense is recalibrated proportional to its relative reliability. The reliability-based model for cue recalibration is a logical extension of optimal cue integration in the sense that reliability directly guides the degree of recalibration for each modality.

### Fixed-ratio cross-modal recalibration

In contradiction to the results just discussed, a recent study investigating the mechanism underlying visual-vestibular recalibration found evidence that each modality is recalibrated in a fixed ratio and, thus, provided evidence against reliability-based cue recalibration [61]. More specifically, after exposing humans and monkeys to a systematic discrepancy between visual and vestibular heading direction, both cues significantly shifted in the direction required to reduce cue conflict, but the amount of recalibration in either modality did not depend on the measured relative reliabilities of the two cues for each participant [61]. A model assuming a fixed ratio between the degree of recalibration of each sense, independent of relative reliability, best captured the data. According to this model, in the event of a cross-modal cue discrepancy, each modality should be adapted towards the other with a fixed, modality-specific recalibration rate, independent of the reliability of each sensory cue.

### Causal-inference and cross-modal recalibration

Both reliability-based and fixed-ratio models of cross-modal recalibration typically assume that the brain acts upon the discrepancy between the two sensory cues. However, two discrepant cues can lead to very different percepts, depending on each cue’s reliability and an inference about whether they have shared or separate origins. Cross-modal recalibration might take this perceptual inference into account rather than relying on the incoming sensory cues alone. This hypothesis makes intuitive sense: cues that are not perceived as coming from a common source should not be used to recalibrate one another. Indeed, causal-inference models of cross-modal recalibration have successfully predicted visual-auditory [62] and visual-tactile [31] ventriloquism aftereffects. Unlike the two linear models of cross-modal recalibration, in this model, recalibration is not based on a mere comparison of the two sensory cues, but rather relates the cues to the perceptual estimate and by doing so incorporates causal inference and cue reliability. Thus, the conflict between previous findings regarding the role of cue reliability for recalibration might reflect different scenarios regarding the perception of the two discrepant sensory signals rather than diverging underlying mechanisms.

### Preview

In this study, we contrasted all three accounts of cross-modal recalibration by fitting the corresponding models to data from an audiovisual ventriloquism-aftereffect study. Across sessions, we manipulated cue reliability, a determinant of reliability-based and causal-inference-driven cross-modal recalibration. Specifically, we selected stimuli for which visual reliability was greater than, approximately equal to, and smaller than auditory reliability. Additionally, we controlled for the effect of modality-specific spatial biases on the spatial perception of the two sensory cues by choosing for each participant visual stimulus locations that matched perceptually with the locations of the auditory stimuli. Each session of the recalibration experiment comprised three phases:

1. baseline: participants localized unimodal visual and auditory stimuli,
2. recalibration: participants were presented with a series of audiovisual stimuli with a constant spatial discrepancy between the auditory and the visual stimulus, and
3. post-recalibration: unimodal localization was remeasured.

With decreased visual reliability, many participants showed either increasing or first increasing and then decreasing auditory recalibration. No clear pattern of visual recalibration was found. These results cannot be explained by either the reliability-based or the fixed-ratio model. The causal-inference model, on the other hand, is able to capture these idiosyncratic variations of recalibration based on individual differences in cue reliability, modality-specific biases, and an a priori assumption of the probability of a common cause. Thus, the model comparison suggests that cross-modal recalibration is driven by a comparison between sensory cues and the corresponding perceptual estimates, which in turn are determined by causal inference.

## Results

### Cue reliability

In the first part of the study, participants’ individual unimodal spatial reliability for one auditory stimulus and three visual stimuli was estimated in four separate sessions using a unimodal spatial-discrimination task. The auditory stimulus was a brief noise burst. Each visual stimulus was a collection of ten Gaussian blobs, the locations of which were randomly sampled from a Gaussian distribution. Visual reliability was manipulated by varying the horizontal spread of the blobs’ locations (example visual stimuli are shown in Fig 1A). The unimodal spatial-discrimination task was carried out using a two-interval forced-choice (2IFC) procedure. In each trial, two stimuli of the same modality were presented sequentially. In one randomly determined interval, participants were presented with a standard stimulus located at the center of the screen. In the other interval, participants were presented with a test stimulus, the location of which was determined by a staircase procedure. Participants were then asked to report which interval contained the stimulus that was located farther to the right. Feedback was provided after the response (Fig 1B). The responses were coded as the probability of reporting the test stimulus to be more rightward than the standard stimulus. For each stimulus condition, we fitted a cumulative Gaussian distribution with a lapse rate (constrained to be less than 6%) to the responses as a function of the distance between the test and standard stimuli (Fig 1C).

**Fig 1.**
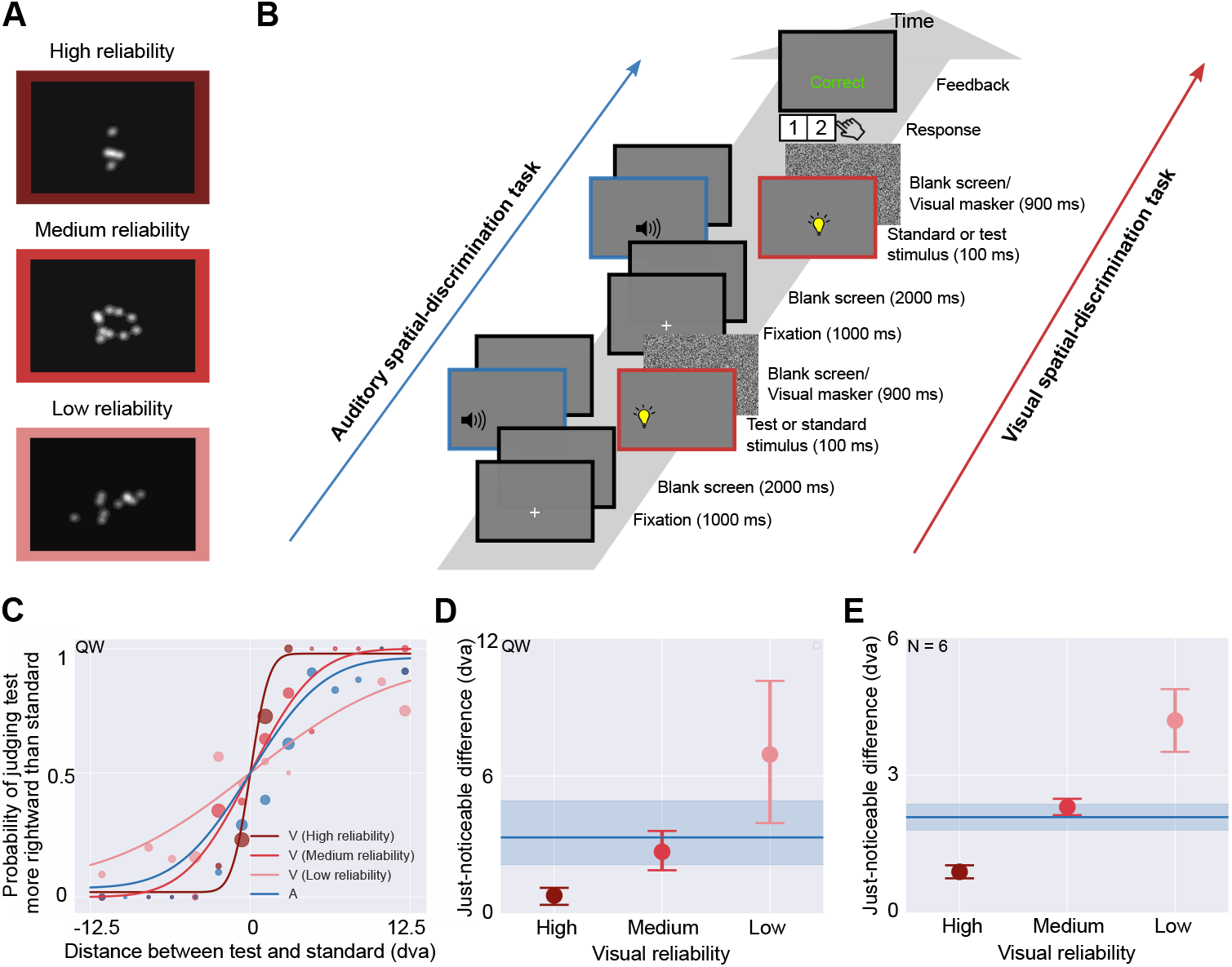
Visual stimuli, experimental procedure, and results for the unimodal spatial-discrimination task. (A) Example stimuli (dark pink: high reliability; pink: medium reliability; light pink: low reliability), visual reliability was manipulated by varying the spread of the locations of the ten Gaussian blobs that constituted a stimulus. Contrast exaggerated for better visibility. (B) Task timing (blue: auditory stimuli; pink: visual stimuli). Participants were presented with a standard stimulus in one interval and a test stimulus in the other interval in random order. After stimulus presentation, they used a keypad to report which interval contained the stimulus located farther to the right. Feedback was provided. (C) Psychometric functions for representative participant QW. The probability of judging the test stimulus to be right of the standard stimulus is plotted as a function of the distance between the two stimuli, with negative numbers indicating that the test stimulus was located to the left of the standard stimulus (presented at straight-ahead). Circular markers: binned psychometric data (bin size = 1.8°). The area of each marker is proportional to the number of trials within the bin. Solid curves: psychometric functions fit to the data. (D) Estimated just-noticeable difference (JND) of each visual stimulus type for QW. Solid line: auditory JND. Error bars and blue area: 95% bootstrapped confidence intervals. (E) Mean JNDs averaged across participants (± SEM).

To compare spatial-discrimination performance across stimulus conditions, we computed the just-noticeable difference (JND; Fig 1D,E), defined as half the distance between test stimulus locations corresponding to probabilities of 0.25 and 0.75 according to the psychometric function (the best-fitting cumulative Gaussian unscaled by lapse rate). Statistical analysis of the JNDs (S1 Appendix: S1.1) revealed that by varying the spread of the collection of blobs, we successfully identified visual stimuli with spatial-discrimination thresholds (i) smaller than, (ii) comparable to, and (iii) larger than the auditory discrimination threshold.

### Modality-specific spatial biases

In the second part of the study, we mapped participants’ modality-specific biases in spatial perception by measuring shifts in visual relative to auditory localization. To this aim, participants completed a bimodal spatial-discrimination task. Auditory stimuli were presented at four different locations (±2.5 and ±7.5° relative to straight-ahead). In each trial, one of the four auditory standard stimuli and a visual test stimulus were presented in random order. The position of the visual test stimulus was controlled by an adaptive procedure. Participants reported whether the visual test stimulus was located to the left or right of the auditory standard stimulus. Feedback was not provided (Fig 2A). Only the high-reliability visual stimulus was used, to measure the biases in as noise-free a manner as possible. The responses were coded as the probability of reporting the visual test stimulus to be located more rightward than the auditory standard stimulus. We fitted a cumulative Gaussian distribution with a lapse rate (constrained to be less than 6%) to the responses as a function of visual test stimulus location. Four separate psychometric functions were fitted, one for each auditory standard stimulus location (Fig 2B). For each psychometric function, we calculated the point of subjective equality (PSE), defined as the distance between the visual test and the auditory standard stimulus that corresponded to a probability of 0.5 according to the psychometric function. A linear regression was used to model the PSEs as a function of auditory location (Fig 2C). The estimated slopes for five out of six participants were significantly larger than 1 (Fig 2D). Thus, the auditory standard stimuli were perceived as shifted towards the periphery relative to the visual test stimuli, which is in line with previous findings ([46, 63–65] but see [66]). Four participants showed significant negative *y*-intercepts, they perceived the auditory stimuli as shifted to the left relative to visual stimuli. From the linear regression line through the PSEs, we extracted the visual stimulus locations that were perceived to be co-located with the four auditory stimulus locations and used them in the subsequent recalibration phase.

**Fig 2.**
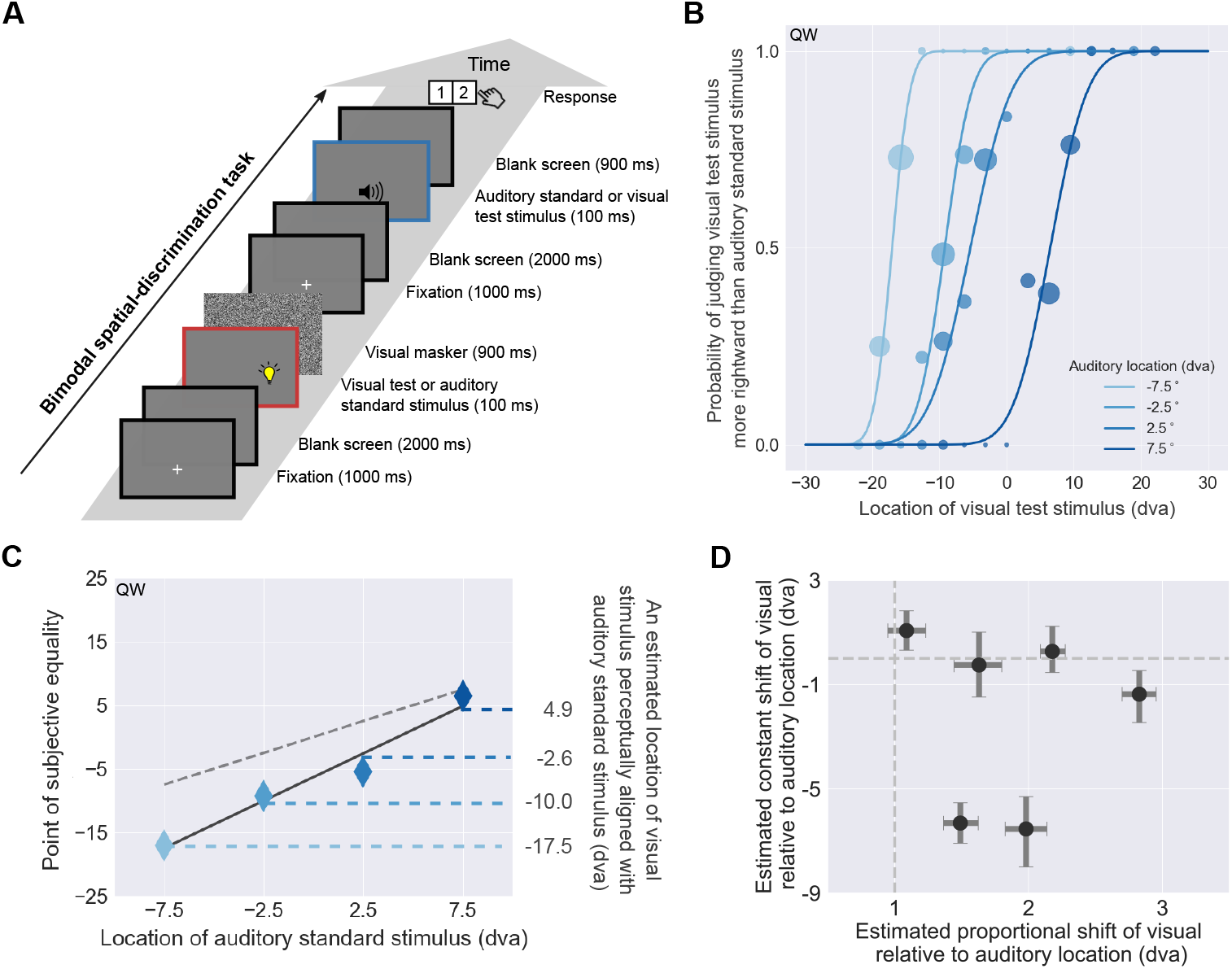
Experimental procedure and results for the bimodal spatial-discrimination task. (A) In each trial, participants were presented with a visual test stimulus (high reliability) in one randomly chosen interval and an auditory standard stimulus in the other interval (four possible auditory locations, ± 2.5 and ±7.5°). After stimulus presentation, participants reported whether the visual test stimulus was to the left or right of the auditory standard stimulus. Feedback was not provided. (B) Psychometric functions for participant QW. Probability of judging the visual test stimulus to the right of the auditory standard stimulus is plotted as a function of the visual stimulus location relative to straight-ahead. Dots: binned psychometric data (bin size = 3°). Curves: psychometric functions fitted separately for the four auditory stimulus locations (shades of blue). The area of each marker is proportional to the number of trials after binning. (C) Point of subjective equality (PSE) as a function of the location of the auditory standard. Diamonds: fitted PSEs; error bars: 95% bootstrapped confidence intervals (smaller than the marker size for this participant). Dashed grey line: identity line; solid line: linear regression line; horizontal dashed lines: visual stimulus locations perceived as co-located with the four standard locations for auditory stimuli based on the regression. (D) Estimated constant, location-independent (*y*-axis) and proportional, location-dependent (*x*-axis) perceptual biases of vision relative to audition. The proportional and constant biases equal the slope and intercept of the linear regression line through the PSEs (see (C)). Error bars: 95% bootstrapped confidence intervals. Vertical and horizontal dashed lines correspond to the absence of proportional and constant perceptual biases of vision relative to audition, respectively.

### Localization response precision

In the next part of the study, we measured participants’ localization noise unrelated to spatial perception (e.g., noise due to location memory noise and the ability to get the response device to indicate the intended location). To this aim, participants performed a localization task with maximally reliable visual stimuli. At the same time, they were familiarized with our custom-made pointing device. In each trial, a white dot was displayed briefly at one of eight locations (evenly spaced from −17.5° to 17.5° relative to straight-ahead). Shortly after stimulus presentation, participants moved a visual cursor controlled by the pointing device to the stimulus location. We assumed that the errors were unbiased and independent of stimulus location (see S1 Appendix: S2 for a more complex model and results of a model comparison) and fitted them with a Gaussian distribution centered at zero to estimate the extent of noise corrupting localization responses. Participants’ spatial perception-unrelated localization noise was 1.85° on average (range 1.55–2.29°).

### Recalibration effects

In the final part of the study, participants completed six recalibration sessions (2 recalibration directions × 3 levels of visual reliability). Each session consisted of three phases: (1) Baseline: in each trial, participants were presented with an auditory stimulus, located at ±2.5 or ±7.5° relative to straight-ahead, or with a visual stimulus, the location of which was determined based on the modality-specific biases measured in the bimodal spatial-discrimination task. After each stimulus presentation, participants localized the stimulus by moving a visual cursor to its location (Fig 3A). (2) Audiovisual recalibration: participants were repeatedly presented with audiovisual stimulus pairs consisting of an auditory and a spatially discrepant visual stimulus (Fig 3B). The spatial discrepancy between the two stimuli was kept constant in perceptual space (Fig 3C). More specifically, in the visual-left-of-auditory condition, the leftmost visual location was paired with the second leftmost of the four standard auditory stimulus locations, the second with the third, and the third with the rightmost auditory stimulus location. For the visual-right-of-auditory condition, it was the other way around. In each trial, participants localized the stimulus of one modality, with the modality being cued after stimulus presentation. (3) Post-recalibration: participants completed the same unimodal localization task as in the baseline phase.

**Fig 3.**
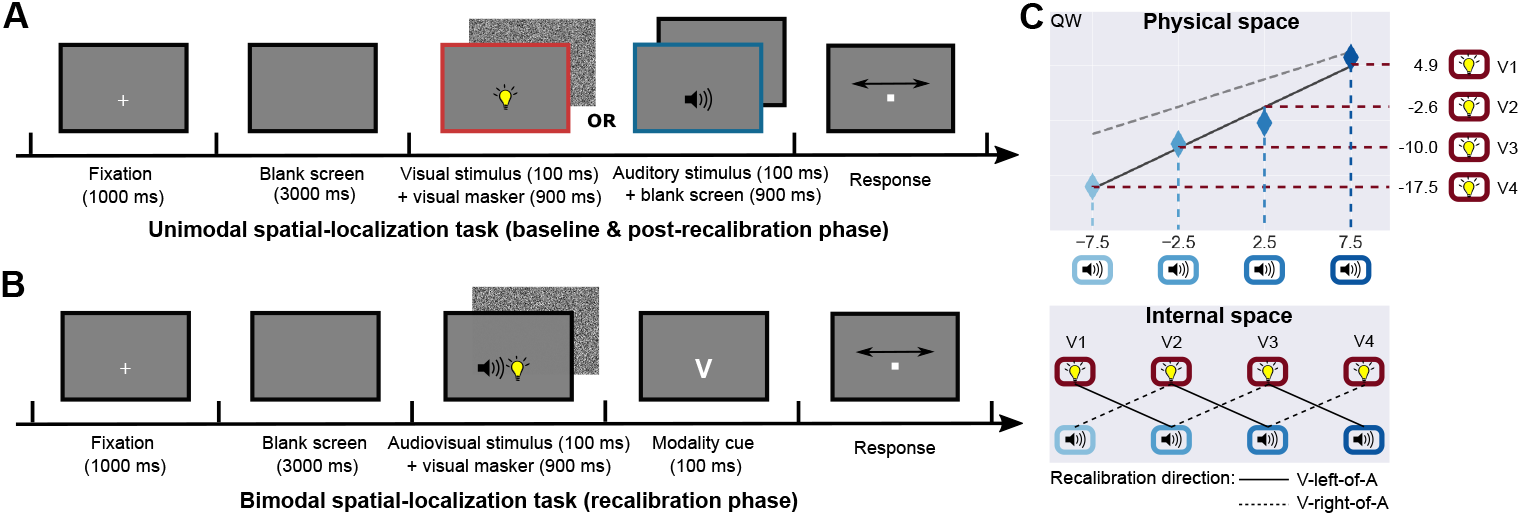
Experimental procedure and conditions for the recalibration sessions. (A) Timeline of the unimodal spatial-localization task conducted during baseline and post-recalibration phases. In each trial, participants were presented with either a unimodal visual or auditory stimulus, and they indicated its location using a visual cursor displayed on the screen. Feedback was not provided. (B) Timeline of the bimodal spatial-localization task conducted during the recalibration phase. In each trial, participants were presented with a spatially discrepant audio-visual stimulus pair, and they were asked to localize one of the modalities, cued after stimulus presentation (V: localize the visual component, A: localize the auditory component). Feedback was not provided. (C) Stimulus locations in physical and perceptual space. Top panel: the physical locations of visual stimuli (*y*-axis) perceptually aligned with four auditory stimuli (*x*-axis) for a representative participant QW; bottom row: the four auditory and visual stimuli presented in both tasks were aligned in internal perceptual space; pairs with a constant discrepancy were presented during the recalibration phase, solid lines: location pairs presented in the visual-left-of-auditory condition; dashed lines: location pairs presented in the visual-right-of-auditory condition.

After repeated exposure to spatially discrepant audiovisual stimuli, in the post-recalibration phase the perceived locations of the auditory stimuli were shifted in the direction of the discrepant visual stimuli presented during the recalibration phase. There was also a shift in the perceived location of visual stimuli, but the magnitude was much smaller compared to the auditory shift (see Fig 4A for an example participant and session).

**Fig 4.**
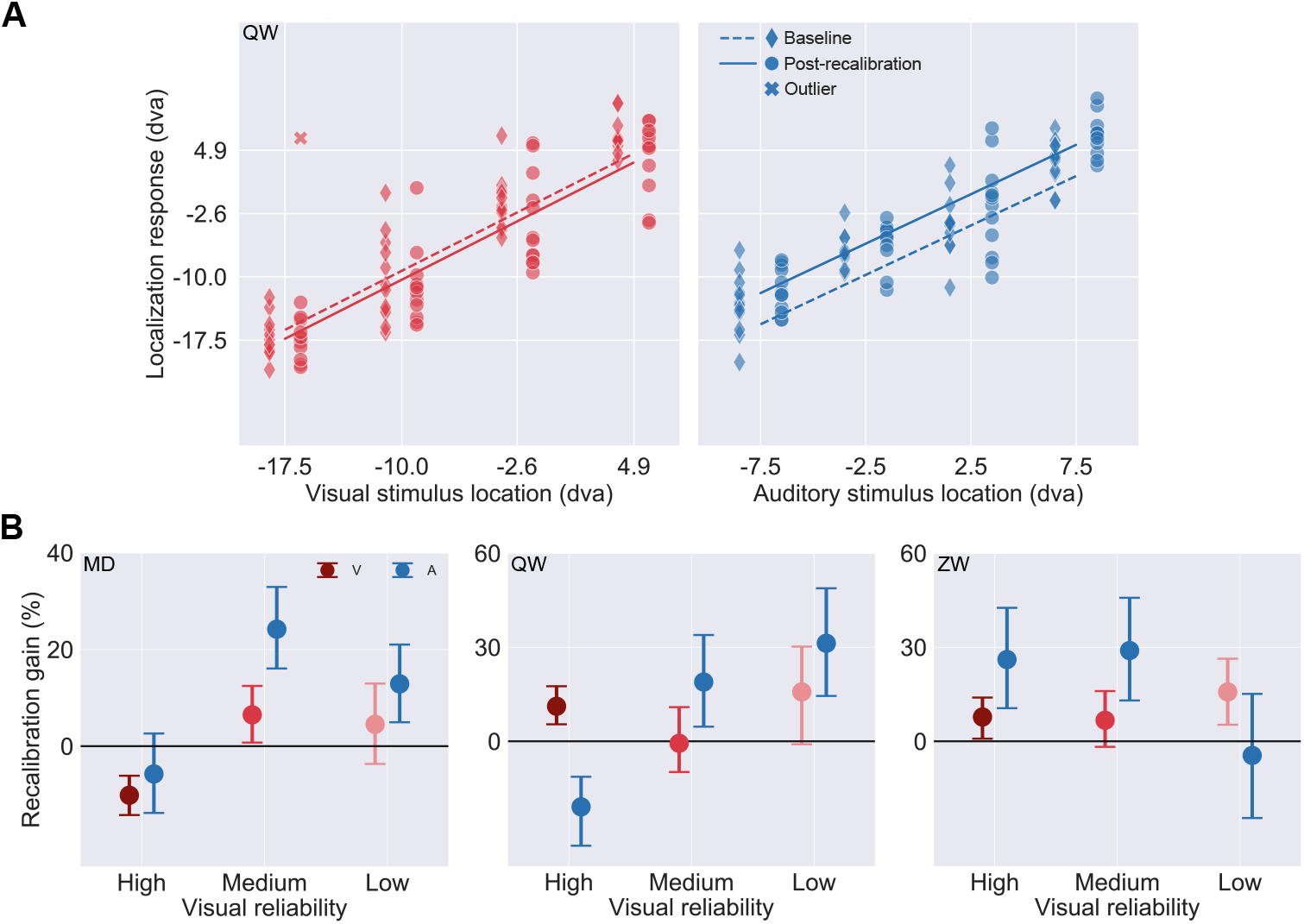
Recalibration effects. (A) Localization responses as a function of stimulus location separately for the baseline (diamonds, jittered to the left) and the post-recalibration phase (circles, jittered to the right) for participant QW in the visual-right-of-auditory and low-visual-reliability condition. Pink: visual localization responses; blue: auditory localization responses; crosses: outlier responses; dashed lines: linear regressions fit to the localization responses in the baseline phase; solid lines: linear regressions fit to the localization responses in the post-recalibration phase. (B) Auditory and visual recalibration gain (shift relative to discrepancy during the recalibration phase) as a function of visual stimulus reliability for three participants. Error bars: 95% bootstrapped confidence intervals.

To examine the amount of recalibration of one modality by the other independent of stimulus location, we fitted the localization responses in both phases as a function of stimulus location using linear regression (Fig 4A). More specifically, for each modality, we fitted one regression line to the localization responses in the baseline phase from all six sessions (the baseline localization responses should not vary across days) and six regression lines to the localization responses in the post-recalibration phase, one for each session (2 recalibration directions × 3 visual reliabilities). We fitted with the constraint that all seven linear regression lines share the same slope, because we assumed that the amount of recalibration does not vary across locations. To quantify the amount of recalibration, we computed the difference of the regression intercepts between the baseline and the post-recalibration phases. To account for the direction of recalibration, shifts from baseline to post-recalibration phase in the direction of the other modality were coded as positive. As no significant effects of recalibration direction on the amount of recalibration emerged in our statistical analysis (S1 Appendix: S1.2), we averaged the amount of recalibration across the two recalibration directions (visual to right/left of auditory) for each level of visual reliability. To normalize the amount of recalibration, we divided the difference by the spatial discrepancy between the auditory-visual stimulus pairs in physical space.

Participants exhibited qualitatively different trends regarding the influence of visual stimulus reliability on the extent of auditory recalibration (Fig 4B for three representative participants). For the majority of participants (three out of six), auditory recalibration gain was a non-monotonic function of visual stimulus reliability. Two participants showed a monotonic increase in auditory recalibration gain with increasing visual stimulus reliability. One participant showed a monotonic decrease in auditory recalibration gain as visual reliability increased.

Given the idiosyncratic and diverse influences of visual stimulus reliability on auditory recalibration gains across participants, it is not surprising that our statistical analysis revealed no significant main effects or interactions involving visual stimulus reliability (S1 Appendix: S1.2). Here, we tested functional models of audiovisual recalibration. A good model of the mechanism governing cross-modal recalibration should be able to capture each participant’s behavior even if participants exhibit qualitatively different effects.

### Recalibration models

To understand the mechanisms of cross-modal recalibration, we compared participants’ behavior to three existing models of cross-modal recalibration: (1) a reliability-based, (2) a fixed-ratio, and (3) a causal-inference model (for details see Models of audiovisual recalibration). These three models differ in their assumptions about the way in which the amount of recalibration for each sense is determined. As a consequence, these three models make different predictions for the influence of visual reliability on audiovisual recalibration.

All three models assume that stimuli lead to noisy and biased sensory measurements. The variability of the measurements is determined by stimulus reliability, with lower reliability meaning more variation in the measurements. To compensate for systematic discrepancies between the senses, each measurement is shifted in a modality-specific manner, and these measurement shifts are constantly updated. Artificially created systematic discrepancies between co-occurring auditory and visual stimuli as induced in the recalibration phase of our study lead to systematic discrepancies (corrupted by noise but still systematic) between auditory and visual measurements. Visual and auditory measurements will be shifted towards the direction of the stimulus discrepancy during the recalibration phase.

#### The reliability-based model

According to this model, the brain determines the amount of recalibration (i.e., the shift updates) for each modality based on the reliabilities of both sensory measurements. The shift for one modality is updated by a fraction of the difference between the two measurements that is proportional to the other modalities’ relative reliability. Across trials, the amount of recalibration depends not only on stimulus reliability, but also on a common learning rate *α* for both modalities. This model predicts increasing visual and decreasing auditory recalibration gains with decreasing visual stimulus reliability.

#### The fixed-ratio model

According to this model, the brain determines the amount of recalibration based on the modalities of both sensory measurements. The measurement shift for each sense is updated by a fraction of the difference between the two measurements. This fraction is determined only by modality-specific learning rates. Therefore, this model predicts modality-specific recalibration gains but no influence of visual stimulus reliability on recalibration gains for either modality.

#### The causal-inference model

According to this model, the brain determines the amount of recalibration for each modality based on the difference between a measurement and the corresponding location estimate. The measurement-shifts are updated by a fraction of this difference, either implemented as modality-specific learning rates (denoted as model CI) or as a supra-modal learning rate (model 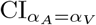). In this model, cross-modal recalibration depends on stimulus reliability, modality-specific localization biases (i.e., current measurement shifts), and inference about a common cause for both measurements by reference to the final location estimates. Cue reliability influences the final location estimates and thus the amount of recalibration in two ways: (1) it affects the integrated and segregated intermediate location estimates, which make up the final location estimate and (2) it affects the likelihood of a common cause given the measurements, which determines the weighting of the two intermediate location estimates. Due to this nonlinear influence of stimulus reliability in combination with participants’ spatial biases and common-cause prior, the model predictions regarding the effects of visual stimulus reliability on recalibration gains are diverse. More specifically, the model can predict monotonically decreasing, monotonically increasing, and non-monotonic recalibration gains, depending on the reliability of the cues, participants’ prior regarding a common cause for vision and audition, and their spatial biases for auditory relative to visual spatial perception.

To fit the free parameters of these models, we minimized the difference between participants’ observed behavior and model predictions for unimodal localization responses in both baseline and post-recalibration phases. We additionally used the data from the unimodal and bimodal spatial-discrimination tasks as well as the pointing practice task to constrain the parameter estimates. Due to the fact that the measurement-shift updates are sequentially dependent and fitting models to such data is extremely time-consuming and computationally expensive, we did not compare participants’ localization responses during the audiovisual recalibration phase with model predictions.

## Modeling results

### Model predictions

The causal-inference model is the only model that can capture the idiosyncratic influence of visual stimulus reliability on auditory recalibration by vision that emerged across participants, the monotonic increase, the monotonic decrease, and the non-monotonic change in auditory recalibration gains with decreasing visual stimulus reliability (Fig 5A). The causal-inference model with a supra-modal learning rate can capture some of the reliability-dependent auditory recalibration gains and outperforms the causal-inference model with modality-specific learning rates for these participants. Neither the reliability-based nor the fixed-ratio model can predict the influence of visual stimulus reliability on auditory recalibration for most participants (Fig 5B; S1 Appendix: S11-S12). The three models do not differ in terms of capturing visual recalibration gains and modality-specific biases (S1 Appendix: S3).

**Fig 5.**
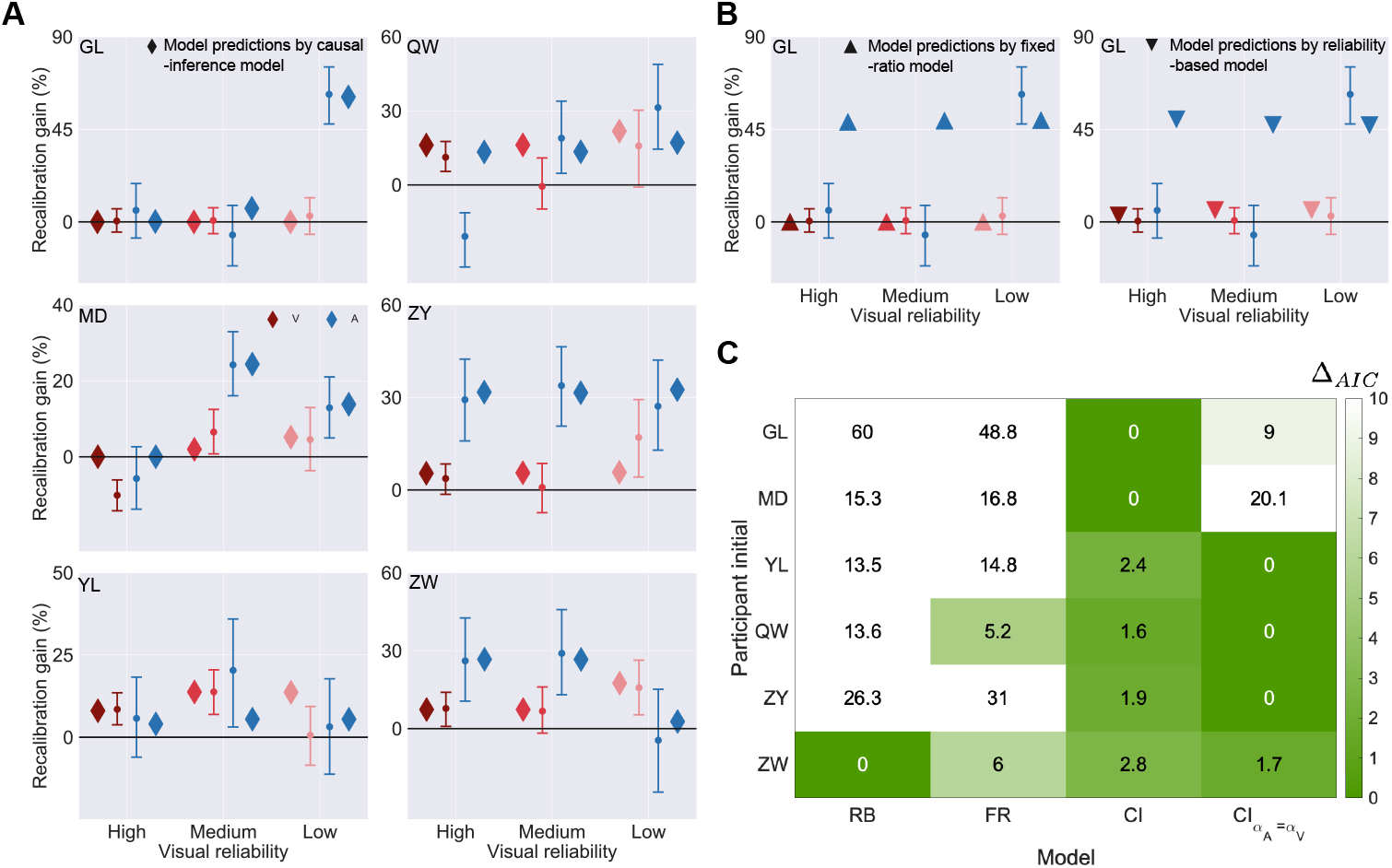
Model predictions and model comparison. (A) Model predictions by the causal-inference model (diamonds) for auditory (blue) and visual (pink) recalibration gains as a function of visual reliability for all participants (panels). Error bars: 95% bootstrapped confidence intervals. (B) Model predictions by the fixed-ratio model (left panel) and the reliability-based model (right panel) for participant GL. (C) Model comparison indices (smaller values indicate more evidence for each model).

### Model comparison

To compare model performance quantitatively, we computed the Akaike Information Criterion (AIC) for all three models [67] and then calculated relative model-comparison scores, Δ_AIC_, which relate the AIC value of the best-fitting model to that of each of the other models (a high delta value indicates stronger evidence for the best-fitting model, i.e., the model with the lowest AIC, and weaker evidence for the comparison model; Fig 5C). Δ_AIC_ values comparing both the reliability-based and the fixed-ratio to the causal-inference models were large for the majority of participants, revealing substantial evidence for the causal-inference model of cross-modal recalibration.

## Discussion

In this study, we investigated the mechanism underlying cross-modal recalibration. To this aim, we measured the effects of visual stimulus reliability, a potential determinant of cross-modal recalibration, on audiovisual spatial recalibration. To induce recalibration, we repeatedly exposed participants to audiovisual pairs with a constant spatial discrepancy; to measure recalibration, we analyzed the localization of auditory and visual stimuli presented unimodally before and after the exposure. Auditory localization was recalibrated by vision, yet, the influence of visual stimulus reliability on auditory recalibration differed in size across participants. To scrutinize the mechanisms of cross-modal recalibration in each participant, we compared their behavior to three models of recalibration: (1) a reliability-based model, which assumes that the amount of cross-modal recalibration depends on the relative reliability of the cues that indicate an inter-sensory conflict, (2) a fixed-ratio model, which assumes that the amount of cross-modal recalibration is fixed, dependent only on the modalities in conflict and thus independent of cue reliability, and (3) a causal-inference model, which ties recalibration to the percept of a cue. This percept depends on the other cue, causal inference of the cues coming from a common source as well as cue reliabilities, and modality-specific spatial biases. Only the causal-inference model captured the idiosyncratic influences of cue reliability on cross-modal recalibration based on individual differences in the determinants of the percept.

### Only the causal-inference model captures the diverse influences of visual reliability on auditory recalibration by vision

Our results demonstrated diverse influences of visual stimulus reliability on auditory recalibration. For half of the participants, auditory recalibration gains were maximal at medium visual stimulus reliability. For some other participants, auditory recalibration gains increased with decreasing visual stimulus reliability. Neither of these patterns can be replicated by models of recalibration that assume the amount of recalibration relies directly on cue reliability [53, 60]. These models can only predict decreasing recalibration with decreasing stimulus reliability of the other modality, as has been found previously [58, 60]. Thus, the best prediction these models could produce for either monotonically or non-monotonically increasing recalibration gain with decreasing stimulus reliability of the other modality was no influence of stimulus reliability (Fig 5B, right panel). The observed recalibration gains are also at odds with models of recalibration that assume the amount of recalibration relies only on the identity of the two modalities in conflict [61, 68]. These models predict no influence of stimulus reliability on recalibration. Crucially, the causal-inference model of cross-modal recalibration [31, 62] captures all the observed influences of stimulus reliability on cross-modal recalibration and is able to replicate all previous patterns of results [60, 61] based on individual differences in cue reliability, the common-cause prior, and modality-specific spatial biases.

It is remarkable that the causal-inference model of cross-modal recalibration is capable of producing qualitatively different patterns of results for the amount of cross-modal recalibration as stimulus reliability changes [31]. Here, for the first time, by fitting this model to the diverse patterns of results we observed, we were able to reveal qualitatively different predictions regarding the amount of recalibration as a result of differences in the combinations of the prior assumption of a common source for both cues and the uncertainties associated with these cues. Additionally, we discovered that biases in the spatial perception of both modalities directly influence the size of the perceived spatial discrepancy and therefore the amount of recalibration. Based on our experience with the model, we next provide an intuition of how these three factors, according to the causal-inference model, influence cross-modal recalibration.

In the models, auditory and visual measurements are biased. After every encounter with an audiovisual stimulus pair, the bias for each modality is updated. Subsequent measurements of both modalities are shifted a little more in the direction of the measurement of the other modality, and these measurement-shifts are updated after each encounter. These updates of the measurement shifts depend on the difference between the measurement and the corresponding location estimate. The recalibration gains are the percentage of the accumulated shifts after completion of the recalibration phase relative to the imposed discrepancy.

#### The role of cue reliability for cross-modal recalibration

In the audiovisual recalibration task, participants were presented with a spatially discrepant audiovisual stimulus pair, resulting in an auditory and a visual measurement in the brain. For each measurement, the brain assigns probabilities to locations in the world based on the measurement and knowledge about its reliability as codified in the likelihood function. When the visual cue is extremely reliable, the likelihood function is narrow and has little overlap with the likelihood function of the auditory location, which is displaced from the visual one due to the spatial discrepancy between the physical stimulus locations (Fig 6A, top panel). The separation between these likelihood functions results in a small likelihood of the auditory and visual stimuli coming from a common source, and hence a low posterior probability of a common source (Fig 6B). This in turn leads to a small difference between the measurement and the final location estimate (Fig 6D) because of the way the final location estimate is calculated. We assumed that participants used a model-averaging strategy to compute their final location estimates. They combine two intermediate estimates, one given a common cause (Fig 6C) and the other given separate causes, each weighted by the posterior probability of the corresponding causal structure. The estimate given separate sources is closer to the measurement than that given a common source as only the latter is dependent on both cues. A small posterior probability of a common source results in final location estimates that are close to the separate-cause estimates, which in turn leads to a small difference between the measurement and the final location estimate (Fig 6D). As the measurement-shift updates are based on the difference between a measurement and the corresponding final estimate, small recalibration gains are predicted when cues are highly reliable.

**Fig 6.**
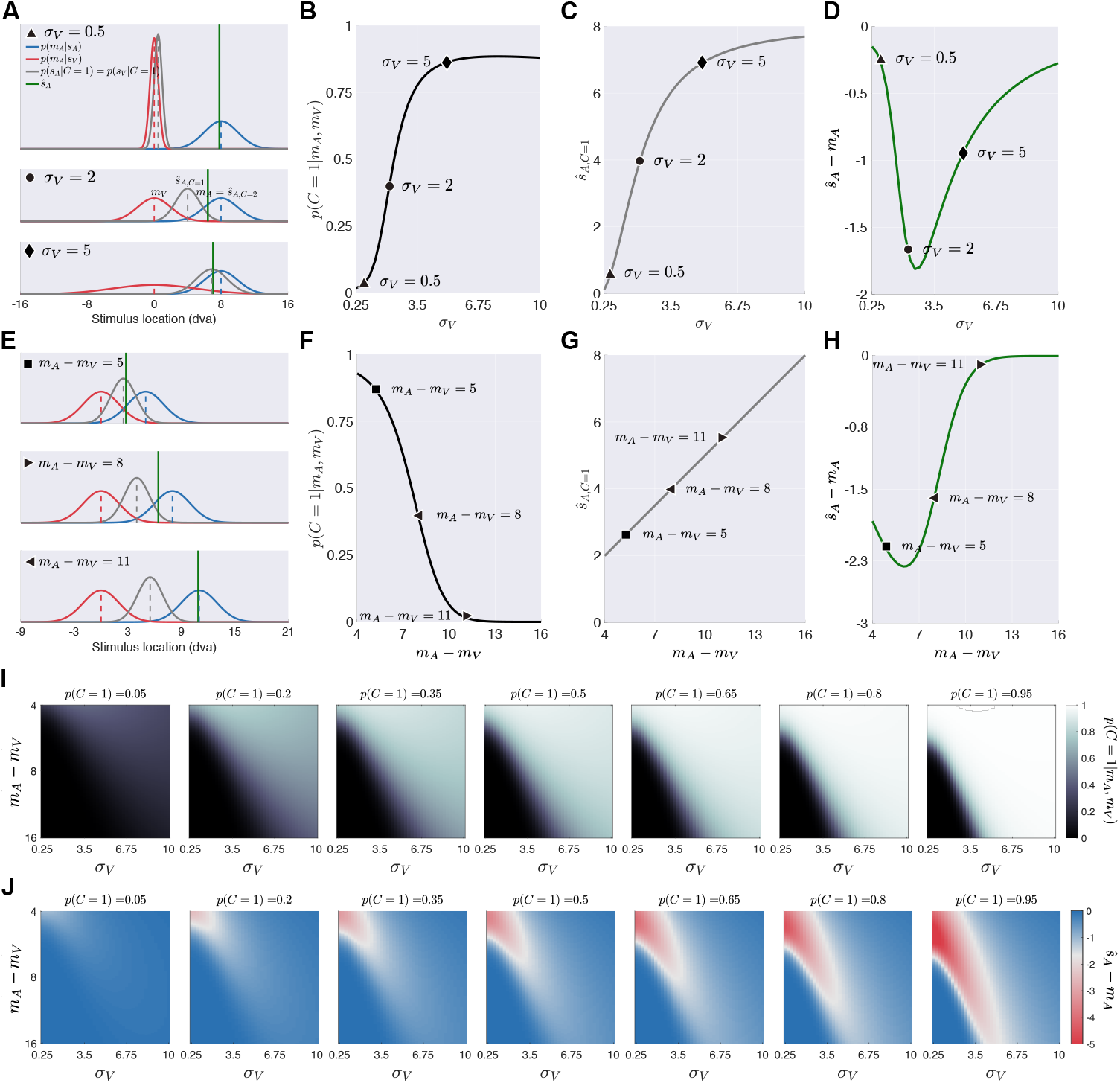
Three factors influencing the degree of recalibration. The effects of three crucial factors, cue reliability, spatial biases and the common-cause prior on three sequential determinants of the degree of cross-modal recalibration: 1) the posterior probability of a common cause *p*(*C* = 1|*m*_*A*_, *m*_*V*_), 2) the auditory location estimate when a common cause is assumed *ŝ*_*A,C*=1_ and 3) the discrepancy between a measurement and the corresponding percept, *ŝ*_*A*_ − *m*_*A*_. (A) The non-linear effect of visual stimulus reliability (triangle: high, i.e., small *σ*_*V*_; circle: medium, i.e., medium *σ*_*V*_; diamond: low, i.e., large *σ*_*V*_) on the final auditory percept *ŝ*_*A*_ (green lines). The visual stimulus is located at the center of the display and the auditory stimulus location is 5° to its right (*s*_*V*_ = 0°, *s*_*A*_ = 5°). The observer has a flat prior over stimulus locations, a common-cause prior of 0.5, and a fixed peripheral bias for audition, so that the auditory measurement (vertical blue dashed lines) is shifted to 8°, no such shift is assumed for the visual measurement (vertical red dashed lines). For simplicity here we assume the noisy measurements are equal to their expected values (0 and 8°). Blue and red curves: likelihood functions of auditory and visual location, respectively. The blue dashed line and curve are also the auditory location estimate and the posterior distribution over auditory locations when separate causes are assumed. Grey dashed lines: the auditory/visual location estimate *ŝ*_*A,C*=1_ (i.e., common cause). Grey curves: posterior distribution of audiovisual locations assuming a common cause. (B-D) The effect of visual reliability on the posterior probability of a common cause, *p*(*C* = 1|*m*_*A*_, *m*_*V*_), the integrated location estimate *ŝ*_*A,C*=1_, and the discrepancy between the auditory measurement and location estimate *ŝ*_*A*_ − *m*_*A*_, which directly sets the amount of recalibration. (E) The effect of a spatial discrepancy between auditory and visual measurements, which is determined by the stimulus locations as well as modality-specific biases (square: small; rightward triangle: medium; leftward triangle: large), on the final auditory perceptual estimate *ŝ*_*A*_. Visual and auditory reliabilities (*σ*_*A*_ = *σ*_*V*_ = 2) and the common cause prior *p*(*C* = 1) (*p*(*C* = 1) = 0.5) are held constant here. (F-H) The effect of spatial discrepancy between the auditory and visual measurements on *p*(*C* = 1|*m*_*A*_, *m*_*V*_), *ŝ*_*A,C*=1_ and *ŝ*_*A*_ − *m*_*A*_. (I-J) The joint effects of visual reliability, spatial discrepancy between auditory and visual measurements and the prior assumption of a common cause (panel) on *p*(*C* = 1|*m*_*A*_, *m*_*V*_) and *ŝ*_*A*_ − *m*_*A*_.

As the spatial uncertainty of the visual stimulus increases, the measurements spread more broadly and the likelihood function of hypothesized visual locations around a single visual measurement becomes wider, which leads to a larger overlap with the auditory likelihood function (Fig 6A, center panel). Consequently, a larger overlap between the two likelihood functions leads to a larger likelihood and hence a larger posterior probability of the auditory and visual stimuli coming from a common source. As a result, the final location estimate will be closer to the integrated location estimate and farther away from the measurement, leading to an increase in the shifts of auditory measurements and finally a greater amount of auditory recalibration by vision.

However, the amount of auditory recalibration does not increase monotonically with decreasing visual reliability. Instead, it drops with further decreases in visual reliability. This is because as the visual stimulus becomes very unreliable, the integrated location estimate is dominated by the auditory measurement (Fig 6A, bottom panel). As a result, the final location estimate will also be close to the auditory measurement, leading to small auditory recalibration gains. In sum, the finding of auditory recalibration gains as a non-monotonic function of visual stimulus reliability can be attributed to the non-monotonic influence of visual reliability on the posterior probability of a common source for both stimuli.

Importantly, the external noisiness of the stimuli and participants’ internal sensory noise determine the effect of stimulus reliability on recalibration gains. A study that uses highly reliable stimuli likely finds increasing auditory recalibration with decreasing visual reliability, as all tested stimuli lead to sensory noise that is lower than the turning point corresponding to the highest recalibration gain (Fig 6D). In contrast, if a study uses rather unreliable stimuli or randomly recruits participants’ whose sensory noise is higher, the reverse pattern would emerge. Thus, different studies can lead to very different results regarding the influence of stimulus reliability on cross-modal recalibration depending on the exact nature of the employed stimuli. Put differently, the contradictory results reported previously [60, 61] can both be explained by the causal-inference model of recalibration.

#### The role of modality-specific biases in spatial perception for cross-modal recalibration

Additionally, the amount of audiovisual recalibration depends on individual biases in the relative perception of auditory and visual stimulus location. Our results showed, in agreement with previous studies, that auditory stimuli are perceived as shifted towards the periphery compared to visual stimuli presented in the same location [64, 65]. However, the size of this location-dependent bias varied across participants. As a consequence, the constant discrepancy between physical stimuli in the recalibration phase leads to variable discrepancies between the visual and auditory measurements across participants. More specifically, for participants who showed a very small peripheral auditory spatial bias, on average the difference between the visual and auditory measurements would not be very different from the physical one (Fig 6E, top panel), while for those participants who showed a relatively strong peripheral bias in auditory perception, the difference between the measurements would be much greater (Fig 6E, bottom panel). For participants who have a stronger peripheral bias in auditory spatial perception, the two likelihood functions, which center around the visual and the biased auditory measurements respectively, would be farther apart (Fig 6E), leading to a smaller posterior probability of a common cause (Fig 6F). Therefore, the final auditory location estimate would be closer to the auditory measurement (Fig 6H), resulting in smaller measurement-shift updates, and smaller auditory recalibration gains.

#### The role of prior assumptions regarding a common cause for cross-modal recalibration

In addition to stimulus reliability and spatial discrepancy between the measurements, the strength of the prior assumption of a common cause also impacts the posterior probability of a common cause (Fig 6I) and hence the distance between the final auditory estimate and the measurement (Fig 6J) and the amount of auditory recalibration. As the prior assumption of a common cause increases, the posterior probability of a common cause also increases, leading the final auditory location estimate to be farther away from the auditory measurement and hence yielding a larger measurement-shift update. The interaction of these three factors and the learning rate in the causal-inference model enables it to capture the idiosyncratic patterns we observed here across participants and those observed in previous studies [60, 61].

### Modality-specific spatial biases

Our models differed from previous causal-inference models of cross-modal perception in that we assumed the existence of modality-specific biases but not of modality-specific priors over the space of possible stimulus locations [31, 41, 64]. Both modality-specific biases and priors can account for differences in the localization of auditory and visual stimuli. Modality-specific priors but not biases exert a stronger influence when visual reliability is reduced. To examine which of the two dominated in our study, we conducted a control experiment examining whether the tendency to perceive visual stimuli as shifted towards the central fixation increased with decreasing visual reliability (S1 Appendix: S4). The results suggested that the observed biases in auditory and visual localization responses were dominated by a fixed bias of the sensory measurements rather than due to modality-specific priors. Thus, to avoid overfitting the model, we omitted modality-specific priors in our models and concentrated on modality-specific biases.

The existence of fixed biases in the relative spatial perception of visual and auditory stimuli seems to be at odds with the concept of cross-modal recalibration. If discrepancies between the senses consistently lead to recalibration, why do these biases persist? Likely, perceptual biases reflect adjustments for sensory discrepancies during an early sensitive period of development as has recently been shown for cross-modal biases in temporal perception [31]. Due to the considerable changes the sensory systems undergo during development, the shifts necessary to align the senses during infancy likely differ considerably from those during adulthood. With the closure of the sensory period during late infancy, it becomes impossible to fully compensate for these differences between the senses. In terms of our model, there appears to be a limitation to the shift updates that is set during early infancy.

Our study differs from previous spatial-recalibration studies in that we adjusted the spatially discrepant audiovisual stimuli presented during the recalibration phase for modality-specific biases in spatial perception. During piloting we presented stimuli with a constant spatial audiovisual discrepancy within physical space during the recalibration phase [47–49, 69], and did not find stable recalibration effects. We attributed this to the observation that spatial biases prevented our participants from perceiving audiovisual stimuli aligned in physical space as coming from a common source. To prevent that from happening, we kept the audiovisual spatial discrepancy consistent in perceptual space during the recalibration phase by finding visual locations that were perceived as co-located with four pre-selected auditory locations. This manipulation was successful in that the participants’ auditory spatial perception was recalibrated by vision. We can only speculate as to whether our study differed from other studies in the extent of spatial biases or in the other factors we identified as determinants of cross-modal recalibration and could compensate for the effects of modality-specific spatial biases, cue reliability and the prior assumption of a common cause for both cues.

### Fitting the causal-inference model

Here, we fitted, for the first time, the causal-inference model of recalibration to observed data. To achieve this, we fitted the outcome of the recalibration process rather than its build-up. The recalibration phase itself is characterized by sequential dependence of the measurement shifts and the lack of a closed-form solution for the model. Fitting such models is computationally infeasible. However, by repeatedly simulating the recalibration phase, we obtained an approximation of the distribution of measurement shifts given a set of parameters. The good match between the observed and predicted data as well as our checks of the fitting procedure (S1 Appendix: S5) confirm the validity of our approach.

One negative consequence of not fitting the recalibration process itself is that parameters reflecting stimulus reliability under bimodal conditions are not directly constrained by the data. Given that previous studies indicated that stimulus reliability differs between bimodal and unimodal stimulus presentation in one [31, 36, 70] or the other [71, 72] direction, we incorporated different reliabilities for unimodal and bimodal presentation conditions. Yet, due to the lack of constraint by data under bimodal presentation conditions, we remain cautious in interpreting the estimated reliabilities.

We additionally remain cautious with respect to the interpretation of other parameter estimates, as we found indications for trade-offs between modality-specific biases and a supra-modal prior over stimulus locations (S1 Appendix: S6.1). Such trade-offs made it impossible to estimate both. Due to the importance of modality-specific spatial biases on recalibration outlined above, we assumed a flat supra-modal prior over stimulus location. As a consequence, the estimated modality-specific biases might have been underestimated and sensory reliabilities might have been overestimated, as there is evidence for the existence of both modality-specific biases and a supra-modal prior centered at fixation [31, 41, 64]. Additionally, we show that the prior probability of a common cause and the modality-specific learning rate trade off, an intuitive result as an increase in either factor can lead to a greater amount of recalibration (S1 Appendix: S6.2).

Importantly, even though the possibility of biases in our parameter estimates exists, the mechanisms outlined at the beginning of the discussion all explain the idiosyncratic influence of visual reliability on auditory recalibration and our conclusion remains valid that causal inference-based percepts regulate cross-modal recalibration.

### Conclusion

This study examined the mechanism underlying cross-modal recalibration. To this aim, we measured audiovisual spatial recalibration while varying visual stimulus reliability. Stimulus reliability has been described as one plausible determinant the brain uses to decide which sensory modality should be recalibrated when there is a cue conflict. We found that visual stimulus reliability influenced auditory recalibration in qualitatively different ways across participants. Neither the reliability-based model nor an alternative model that assumes a fixed degree of recalibration for each modality and completely ignores stimulus reliability, could replicate the data. Yet, a causal-inference model was able to capture all the observed diverse influences of reliability on recalibration, including two patterns found in previous, apparently conflicting studies. In this model, recalibration is not based on a mere comparison of the two sensory cues, but rather relates each cue to its corresponding final perceptual estimate, and by doing so incorporates causal inference of a common source for a cross-modal stimulus pair as well as cue reliability and modality-specific perceptual biases into cross-modal recalibration.

## Materials and methods

### Participants

Six participants (three females, aged 22-29, mean: 25, six right-handed), recruited from New York University and naive to the purpose of the study, participated in the experiment. All stated they were free of visual, auditory, or motor impairments. The data of one additional participant (female, aged 21, ambidextrous) were excluded from data analysis and model fitting due to conflicting modality-specific relative biases found in the bimodal spatial-discrimination task and the baseline phase (S1 Appendix: S7). Experimental protocols were approved by the Institutional Review Board at New York University. All participants gave informed consent prior to the beginning of the experiment and five of them were compensated $10 per hour for participation.

### Apparatus and stimuli

The experiment was conducted in a dark and semi sound-attenuated room. Participants were seated 1 m from an acoustically transparent, white screen (1.36 × 1.02 m, 68 × 52° visual angle). An LCD projector (Hitachi CP-X3010N, 1024 × 768 pixels, 60 Hz) was mounted above and behind participants to project visual stimuli on the screen. The visual stimuli were clusters of 10 randomly placed low-contrast (36.55 cd/m^2^) Gaussian blobs (SD: 3.6°) added to a grey background (29.33 cd/m^2^). Blob locations were drawn from a two-dimensional Gaussian distribution (vertical SD: *σ*_*y*_ = 5.4°, horizontal SD: (1) *σ*_*x*_ = 1.1° for the high-reliability condition, (2) *σ*_*x*_ = 5.4° for the medium-reliability condition, and (3) *σ*_*x*_ = 8.7° for the low-reliability condition). We used the centroid of the cluster (rather than the center parameter of the two-dimensional Gaussian used for cluster generation) as the visual stimulus location. Each visual stimulus was presented for 100 ms (6 frames), and followed by a backward masker to erase any visual memory of the stimulus. The backward masker consisted of 10 different frames of visual white noise (randomly black or white checks of 4 × 4 pixels filling the screen), and these 10 frames were presented repeatedly for 900 ms (54 frames).

Behind the screen, there was a loudspeaker (20W, 4ohm Full-Range Speaker, Adafruit, New York) mounted on a sledge attached to a linear rail (1.5 m long). The rail was hung from the ceiling using elastic ropes; it was located 23 cm above the table and 5 cm behind the screen, perpendicular to the line of sight. The position of the sledge on the rail was controlled by a microcomputer (Arduino Mega 2560; Arduino, Somerville, MA, USA). The microcomputer controlled a stepper motor that rotated a threaded rod (OpenBuilds, www.openbuildspartstore.com), which allowed for horizontal movement of the speaker. This way the speaker was moved to its predetermined location before the auditory stimulus was presented. The auditory stimulus was a 100 ms broadband noise burst (0-20.05 kHz, 60 dB), windowed using the first 100 ms (the positive part) of a sine wave with a period of 200 ms.

We were concerned that participants might infer the position of the speaker from the sound that occurred when the speaker moved. We tried to foil this strategy by playing a masking sound during each movement of the speaker; the masking sound was played from an additional loudspeaker, positioned just behind the center of the screen. The masking sound (55 dB) consisted of a recording of the sound generated by a randomly chosen speaker movement plus white noise. To further foil use of auditory cues from the moving sledge, every time the speaker moved from its last position, it first moved to a stopover location before it moved to the target location. The stopover was randomly chosen under the constraint that the total distance the speaker moved and the amount of time the movement took were approximately equal across trials. We carried out a control experiment, which indicated that participants could not infer the speaker position based on speaker movements alone (S1 Appendix: S8).

Responses were given using a numeric keypad in spatial-discrimination tasks, and a pointing device in localization experiments. The pointing device was custom built using a potentiometer (Uxcell 10K OHM Linear Taper Rotary Potentiometer) with a plastic ruler (5 × 17 cm) securely fixed perpendicular to the shaft of the potentiometer. Participants were instructed to place their hands on either side of the ruler and rotate it so that it pointed at the perceived location of the stimulus. A visual cursor (an 8 × 8 pixel white square) was displayed to indicate the selected location. The pointing device was covered by a black box (42 × 30 × 15 cm) so that participants could not see their hands while using the device. A foot pedal was placed on the floor and used to confirm the current position of the pointing device as the response.

Stimulus presentation, speaker movement, and data collection were controlled by a laptop PC running MATLAB R2017b (MathWorks, Natick, MA, USA). Visual stimuli were presented using the Psychophysics Toolbox [73–75].

## Design

### Unimodal spatial-discrimination task

We quantified the reliability of the auditory stimulus and the three visual stimuli for each participant using a unimodal spatial-discrimination task. The visual stimuli had been designed to achieve three distinct levels of reliability for discriminating their spatial locations (Fig 1A). This task allowed us to check whether the auditory stimulus reliability was approximately equal to the medium level of visual stimulus reliability as desired.

### Bimodal spatial-discrimination task

We measured participants’ relative modality-specific localization biases using a bimodal spatial-discrimination task. The aim was to find locations for the visual stimuli that were perceived to be co-located with auditory stimuli at four different locations. These visual stimulus locations were then used in the subsequent recalibration sessions.

### Pointing practice task

We used a unimodal spatial-localization task with maximally reliable visual stimuli to measure additional noise components (e.g., location memory and handling of the pointing device) that affected the precision of participants’ localization responses but were not directly related to spatial perception.

### Audiovisual recalibration tasks

We measured the influence of visual stimulus reliability on audiovisual ventriloquism aftereffects. Three levels of visual stimulus reliability were paired with two recalibration directions, resulting in a total of six conditions, each of which was tested in a separate session. Each session comprised three phases. (1) Baseline: participants localized unimodal auditory and visual stimuli. (2) Recalibration: participants were repeatedly exposed to spatially discrepant bimodal stimuli. They localized the stimulus of one modality, which was cued after stimulus presentation. Three audiovisual stimulus pairs were chosen such that in the visual-to-the-left condition, the leftmost of the visual locations identified in the bimodal spatial-discrimination task was paired with the second leftmost of the four auditory locations, the second with the third, and the third with the rightmost auditory location. In the visual-to-the-right condition, it was the other way around. (3) Post-recalibration: this task was identical to the baseline; ventriloquism aftereffects were measured by comparing localization responses between baseline and post-recalibration phases.

## Procedure

### Unimodal spatial-discrimination task

Participants’ spatial-discrimination thresholds were measured using a 2IFC procedure. In each trial, participants were presented with a standard stimulus in one of two intervals and a test stimulus in the other interval. The order of the standard and the test stimulus was randomized. At the beginning of each trial, a fixation cross was presented at the center of the screen for 1,000 ms, followed by a blank screen presented for 2,000 ms, and then the first stimulus occurred. The stimulus presentation lasted 1,000 ms; the actual stimulus only lasted 100 ms, followed by either a 900 ms backward masker (for unimodal visual stimuli) or a blank screen for 900 ms (for unimodal auditory stimuli). After the first stimulus presentation, the same events (fixation, blank screen, stimulus) were repeated, resulting in a trial duration of more than 8,000 ms. The long pre-stimulus periods were needed to move the loudspeaker to its new position. After the second stimulus period was over, a response probe was displayed and participants indicated by button press which interval contained the stimulus that was farther to the right (Fig 1B). Visual feedback was provided immediately after the response was given (duration: 500 ms). The inter-trial interval was 500 ms.

The standard stimulus was always located at the center of the screen (straight ahead); the location of the test stimulus was controlled by four interleaved staircases, two of which had the test stimulus start at 12.5° (to the right of straight-ahead), and the other two at −12.5° (12.5° to the left of straight-ahead). For the two staircases starting at one side, one followed the two-down-one-up rule (with down being defined as moving the test stimulus leftwards and up as moving rightwards) converging to a probability of 71% [76] of perceiving the test stimulus as farther to the right than the standard stimulus; the other staircase followed the one-down-two-up rule converging to a probability of 29% of choosing the test stimulus as farther to the right than the standard stimulus. The initial staircase step size was 1.9°. It was decreased to 1.0° at the first staircase reversal and again to 0.5° after the third reversal. Each staircase consisted of 40 trials, resulting in a total of 160 trials. To improve the estimation of the lapse rate, we inserted “catch trials”—a test stimulus located distant from the center (±12.5°)—once every 10 trials. A total of 176 trials was evenly split into 4 blocks. Participants completed the spatial-discrimination task for each of the three levels of visual stimulus reliability in random order, and the auditory stimulus was always tested last. Usually participants took about an hour to complete the entire task for one stimulus condition; typically, they completed the testing of all four stimulus conditions in two days.

### Bimodal spatial-discrimination task

Participants’ biases in auditory relative to visual spatial perception were measured using a 2IFC procedure with a standard and a test stimulus presented in random order. The timing of this task was the same as in the unimodal spatial-discrimination task. Participants indicated by button press whether the test stimulus was located to the left or right of the standard stimulus. No feedback was provided (Fig 2A).

The standard stimulus was an auditory stimulus presented at one of four locations (±2.5 or ±7.5°); the test stimulus was the high-reliability visual stimulus used in the unimodal spatial-discrimination task (Fig 1A, first panel). The location of the visual test stimulus was controlled by eight interleaved staircases, two for each of the four auditory locations. Of the two staircases per auditory location, one started the test stimulus at 15° relative to straight-ahead, and the other one at −15°. The staircase with the visual test stimulus starting from the right of the auditory standard stimulus followed the one-down-two-up rule, converging to a probability of 29% of choosing the test stimulus as farther to the right. The staircase with the visual test stimulus starting from the left of the auditory standard stimulus followed the two-down-one-up rule, converging to a probability of 71% of choosing the test stimulus as farther to the right. Staircase step size was updated based on the same rule as described above. Each staircase consisted of 36 trials. A “catch trial” with the visual stimulus being located at ±15° relative to the auditory standard stimulus was inserted once every nine trials, resulting in a total of 320 trials. The session was divided into six blocks. Usually, participants took about two hours to complete all trials.

### Pointing practice task

Participants’ localization precision independent of spatial perception was measured using a localization task with visual stimuli of maximal spatial reliability. In each trial, a white square (8 × 8 pixels ≈0.6° × 0.6°) was displayed on the screen for 100 ms. The stimulus was followed by a 900 ms-long backward masker and 500 ms of blank screen. Subsequently, the response cursor, a green square with the same size as the white square, appeared on the screen. Participants used the pointing device to move the cursor to the stimulus position. Response times were unrestricted. Visual feedback of the cursor location was provided during adjustment, but error feedback was not provided. There were eight possible horizontal positions for the stimulus, evenly spaced from −17.5 to 17.5° in steps of 5°. Each stimulus location was visited 30 times in random order, resulting in a total of 240 trials. The inter-trial interval was 500 ms. This experiment usually took 30 minutes to complete and was typically administered after the bimodal spatial-discrimination task on the same day.

### Audiovisual recalibration task

Participants’ baseline and post-recalibration spatial perception were measured using a unimodal spatial-localization task. In each trial, a fixation cross was presented at the center of the screen for 1,000 ms, followed by a blank screen presented for 2,000 ms, and then either an auditory or a visual stimulus. The inter-trial interval was 100 ms. The auditory stimulus was presented at one of four locations (±2.5 or ±7.5° relative to straight-ahead), and the visual stimulus was presented at the locations that were identified as perceptually co-located with those four auditory locations as estimated in the bimodal spatial-discrimination task for each participant. As in the pointing practice task, participants responded by moving a visual cursor to the stimulus location using the pointing device and confirmed the final position with the foot pedal (Fig 3A). There was no time limit for the response. The location of the visual cursor was provided during the adjustment, but feedback about the localization error was not provided. Each of the four target locations per modality was tested 12 times, resulting in a total of 96 trials administered in pseudorandom order. These trials were split into four blocks. Usually participants took about 25 min to complete all 96 trials.

During the audiovisual recalibration phase, participants were presented with temporally synchronous but spatially discrepant audiovisual stimuli. To ensure that participants were attentive to the stimuli, we asked them to localize either the auditory or the visual component, with the localization modality being cued after stimulus presentation (Fig 3B). The timing of trial events and the response settings were identical to the unimodal spatial-localization task. Each of the three pairs of an auditory and a visual stimulus was repeated 40 times in random order, resulting in a total of 120 trials. These trials were split into four blocks. Usually, participants took about 30 min to complete all 120 trials. Participants first completed the baseline phase, then the recalibration phase, and finally the post-recalibration phase, resulting in a total duration of 80 min for each of the six recalibration sessions.

## Data analysis

### Unimodal spatial-discrimination task

For each session of the unimodal spatial-discrimination task (one auditory and three visual stimulus reliabilities), the data were coded as the probability of identifying the test stimulus as located farther to the right than the standard stimulus as a function of test stimulus location (Fig 1C). We fitted a cumulative Gaussian distribution centered at 0 and with a lapse rate, constrained to be less than or equal to 6% [77], to these psychometric data. The JND was defined as half the distance between the stimulus locations corresponding to a probability of 0.25 and 0.75 of test-stimulus-to-the-right judgments on the best-fitting cumulative Gaussian distribution (unscaled by the lapse rate). We estimated error bars using a bootstrap procedure: we randomly resampled the psychometric data with replacement, creating 1,000 bootstrapped datasets. We re-fitted psychometric functions to each resampled dataset, and took the 2.5 and 97.5 percentiles of the ordered 1,000 JNDs as the bootstrapped confidence interval.

### Bimodal spatial-discrimination task

The data from the bimodal spatial-discrimination task were coded as the probability of identifying the visual test stimulus as located to the right of the auditory standard stimulus as a function of the visual stimulus location. We fitted a cumulative Gaussian distribution to these data, separately for each location of the auditory standard stimulus. Again, we included a lapse rate, constrained to be less than or equal to 6% [77]. The point of subjective equality (PSE) was defined as the location of the visual test stimulus corresponding to a probability of 0.5 for visual-to-the-right-of-auditory responses on the psychometric function. We obtained 95% bootstrapped confidence intervals for each of the four PSE’s (one for each auditory stimulus location) using the same bootstrapping procedure as before. Then, we fitted a linear regression to the four PSE’s of each bootstrap run and from these we obtained 95% confidence intervals for the slope and the intercept of the linear regression. From the linear regression of the original PSEs, we computed the locations of the visual stimulus perceived as co-located with the four auditory locations (±2.5 and ±7.5° relative to straight-ahead). In the subsequent unimodal and bimodal spatial-localization tasks, we presented visual stimuli at the locations obtained from the regression rather than those directly indicated by the PSEs to reduce the effects of random noise during the bimodal spatial-discrimination task.

### Pointing practice task

Data from the pointing practice task (unimodal localization task with maximal-reliability visual stimuli) were first filtered before being analyzed. More specifically, for each stimulus location, we *z*-transformed the data by subtracting the mean from individual localization responses and then dividing by the standard deviation of the demeaned responses. Localization responses with *z*-score *<* −3 or *>* 3 were identified as outliers (range: 0-1.67%). To compute the localization noise unrelated to spatial perception, we fitted a simple model, which assumes location-independent noise and absence of bias, to non-outlier localization responses by a maximum-likelihood procedure (see S1 Appendix: S2 for comparison with a more complex model).

### Audiovisual recalibration task

Data from the unimodal localization tasks were also filtered first before being analyzed. We identified outliers (range: 0.78-1.82%) separately for each modality (visual and auditory) and each experimental phase (baseline, post-recalibration). To this aim, we z-transformed the data in two steps as we did for the pointing practice task. Auditory and visual localization responses in the baseline phase were pooled across all six sessions for computing the mean of localization responses, as the day of testing should not influence localization performance. In contrast, the mean of auditory and visual localization responses in the post-recalibration phase were calculated separately for each of the six conditions as each recalibration condition should influence localization means differently. Then, we computed the standard deviation of the demeaned auditory localization responses pooled across all six session and both phases. The standard deviation of demeaned visual localization responses was calculated separately for each visual-reliability condition given that stimulus reliability should influence localization precision. Localization responses identified as outliers (*z*-scores *<* −3 or *>* 3) were excluded from all further analysis.

To summarize localization responses independent of stimulus location, we fitted linear regression lines to the localization responses as a function of stimulus location, separately for each stimulus modality and each experimental phase (baseline, post-recalibration). For each modality, localization responses in the baseline phase were pooled across all six sessions, as visual reliability should influence the precision but not the accuracy of the localization responses (S1 Appendix: S3). Responses from the post-recalibration phase were fitted separately for each condition, because each recalibration condition should influence localization differently. The seven regression lines per modality were fit with the constraint that all have the same slope.

The amount of recalibration of one modality by the other was defined as the distance between the constant offsets between the baseline and post-recalibration phases (i.e., between the intercepts of the regression lines fit to the baseline and the post-recalibration localization responses). The amount of recalibration was coded as positive if localization responses in the post-recalibration phase were located closer to the location of stimuli in the other modality during the recalibration phase. We calculated the amount of recalibration for each modality and for each recalibration condition (3 visual reliabilities ×2 recalibration directions). We computed the average amount of recalibration across the two recalibration directions for each level of visual reliability because statistical analysis showed no significant main effect of recalibration direction on the amount of recalibration (S1 Appendix: S1.2). Finally, we calculated a unitless recalibration gain by dividing the average amount of recalibration per condition by the spatial discrepancy in physical space during the recalibration phase (5°). To derive confidence intervals, localization responses were resampled, separately for each location, condition (recalibration phase × visual stimulus reliability × recalibration direction), and stimulus modality. Data from the bimodal spatial-localization task conducted during the recalibration phase were not analyzed. Data analysis was done using Python 3.7, R 4.0.2, and MATLAB 2019a.

## Models of audiovisual recalibration

To calibrate the sensory systems, the brain uses discrepancies between information arriving through different senses. However, in the absence of objective information about the calibration status of each sensory system, it remains obscure how the amount of necessary recalibration for each sense is determined. We aimed to distinguish between three models of cross-modal recalibration: (1) recalibration based on stimulus reliability, (2) recalibration based on attributing the measured sensory discrepancy between the senses in conflict in a fixed ratio to each sensory modality, and (3) recalibration based on intra-modal comparisons between sensory measurements and the corresponding perceptual estimates, which in turn rely on causal inference about whether those measurements were caused by a common source. We begin this section by laying out the definition of recalibration underlying all models reported here and then describe the recalibration process during the recalibration phase of our study according to each of the three models. Finally, we provide a formalization of each of the tasks used to constrain the model parameters followed by the details of how models were fit to the data.

### Definition of recalibration

Each stimulus at location *s* in the world leads to a sensory measurement *m*′ in an observer’s brain. This measurement is corrupted by Gaussian-distributed sensory noise. Thus, with repeated presentations of stimuli at location *s*, the sensory measurements correspond to scattered spatial locations *m*′ ~ 𝒩(*s*′, *σ*′^2^). The variability of the measurements is determined by the stimulus reliability 1*/σ*^*′2*^. To allow integration of information from different modalities, measurements from the different modalities are remapped into a common internal reference frame. Hence, the measurement distribution is centered on *s*′, the remapped location of *s*. As part of the remapping process, spatial discrepancies between co-occurring cross-modal stimulus pairs are accounted for by shifting the measurements by a modality-specific amount Δ towards the measurement of the other modality. We model recalibration as the process of updating these shifts after each time the observer encounters a cross-modal stimulus pair [31, 62]. The shift-updates are based on the measurements and therefore stochastic. In the audiovisual recalibration phase in our study, we probed this process by presenting stimulus pairs with a consistent discrepancy. That is, we probed the process of correcting for internal misalignments between the senses by artificially creating a discrepancy between visual and auditory measurements through consistently misaligning the physical stimuli.

We assumed that observers were already calibrated at the beginning of each experimental session. In other words, the remapping included modality-specific shifts that compensated for constant, real-life discrepancies between vision and audition. We use the variables 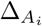 and 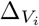 to exclusively capture the change in measurement-shifts due to the spatially discrepant audiovisual stimulus pairs with visual reliability *i* (*i* ∈{1, 2, 3}) presented during the recalibration phase, and we set the modality-specific shifts in the first trial to zero, 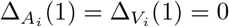. In addition, we assumed that the final shifts after 120 recalibration trials 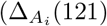 and 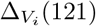) were fully maintained throughout the subsequent post-recalibration phase as observers were not exposed to audiovisual stimulus pairs after the recalibration phase.

We further assumed that stimulus reliability differed between unimodal and bimodal presentations. Note that we denote the visual-reliability condition for variables associated with the auditory modality (i.e., with a subscript *A*_*i*_) when the value of that variable can be impacted by visual measurements (e.g., shifts 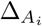 or, below, sensory estimates 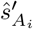), but not otherwise (e.g., measurements 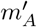 or measurement variances 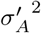 and 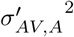).

## Models of the recalibration process

### The reliability-based model of cross-modal recalibration

According to this model, each modality should be recalibrated in the direction of the other discrepant modality by an amount that is proportional to the other modality’s relative reliability [53]. In other words, after every trial, the measurement-shifts, 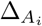 and 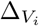, are updated in the direction of the discrepancy between the visual and auditory measurements by an amount proportional to the two modalities’ relative reliabilities as follows:

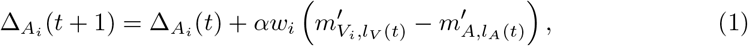

Where

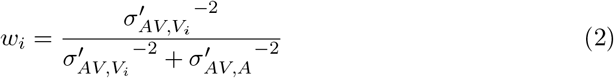

and analogously

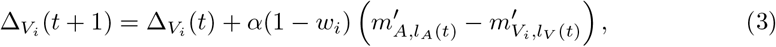

where *l*_*A*_ and *l*_*V*_ index the auditory and visual locations (*l*_*A*_, *l*_*V*_ ∈{1, 2, 3, 4}), *α* denotes a supra-modal learning rate, and *t* denotes trial number.

### The fixed-ratio model of cross-modal recalibration

According to this model, after every trial, 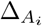 and 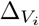 are updated in the direction of the discrepancy between the visual and auditory measurements by a fixed ratio of this discrepancy. The ratio of the update depends solely on the identity of the modality and thus is independent of stimulus reliability [61]. 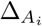 and 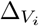 are updated according to the following equations:

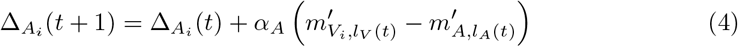

and

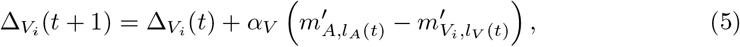

where *α*_*A*_ and *α*_*V*_ are modality-specific learning rates.

### The causal-inference model of recalibration

In this model, the amount of recalibration (i.e., the size of the shift-updates) is directly determined by the discrepancy between a measurement and the corresponding perceptual estimate, 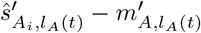 and 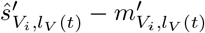 [62]. However, the spatial discrepancy between auditory and visual measurements and the relative reliabilities of both stimuli have indirect influence on the amount of recalibration by means of their influence on the location estimates, 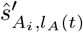 and 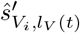. Additionally, the location estimates are contingent on the degree to which the brain infers a common cause or separate causes for the two measurements [29, 62].

The location estimates are a mixture of two conditional location estimates, one for each causal scenario (common source, *C* = 1, or different sources, *C* = 2). In the case of a common source, the location estimate of the audiovisual stimulus pair, 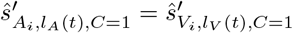, equals the reliability-weighted average of the measurements 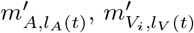 and the mean 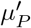 of an internal, Gaussian-shaped, supra-modal prior across stimulus locations with variance 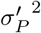 :

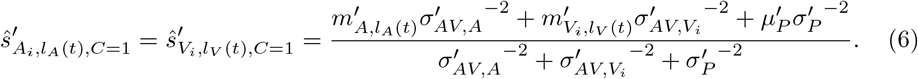

In the case of two separate causes, the location estimates of the auditory and the visual stimulus, 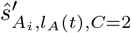 and 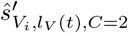, are equal to the reliability-weighted averages of 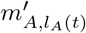 and 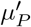 for the auditory estimate, and 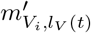 and 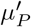 for the visual estimate, respectively:

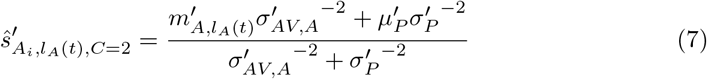

and

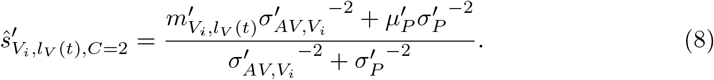

The final location estimates are derived by model averaging. Specifically, the final location estimate 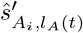 is the average of the conditional location estimates, 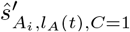 and 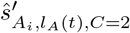, with each estimate weighted by the posterior probability of its causal structure:

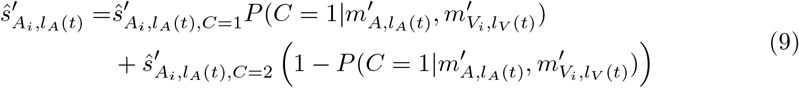

and analogously for the visual location estimate:

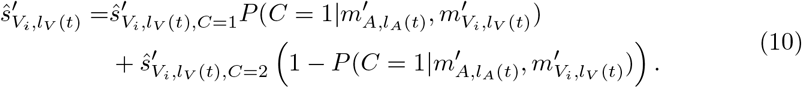

The posterior probability of a common source for the auditory and visual measurements in trial *t*, 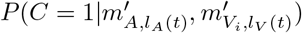, is proportional to the product of the likelihood of a common source for these measurements in trial *t* and the prior probability of a common source for visual and auditory measurements in general, *P* (*C* = 1):

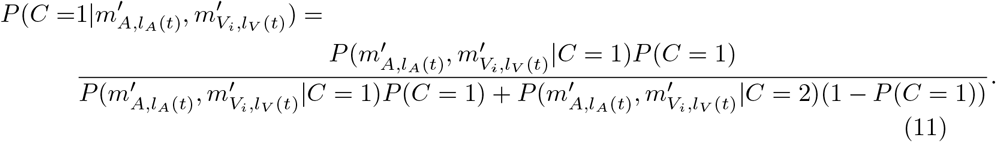

The posterior probability of two separate sources, one for the auditory and one for the visual measurement, is 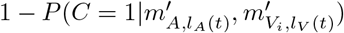.

The likelihood of a common source of the visual and auditory measurements in trial *t*, 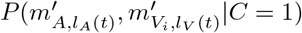, is the product of the likelihood of the internally represented audiovisual stimulus location 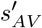 given the auditory measurement, 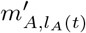, and the visual measurement 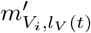, and the supra-modal prior, integrated over all possible remapped audiovisual stimulus locations 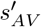 [29]:

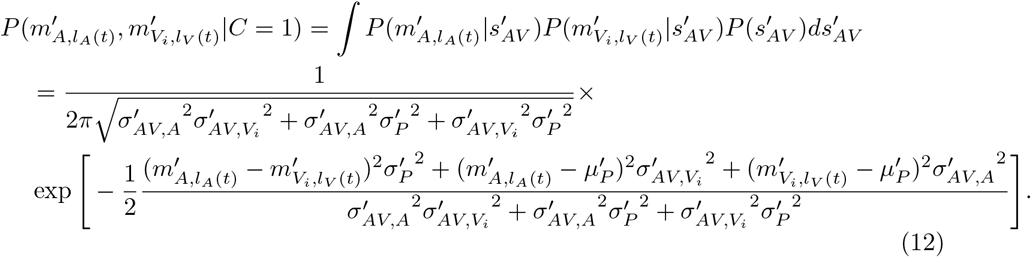

The likelihood of different sources for the visual and auditory measurements in trial *t*, 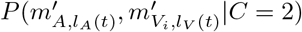 is the product of the likelihood of internally represented auditory and visual stimulus locations 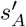 and 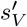 given the auditory measurement, 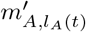, and the visual measurement 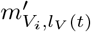, and the supra-modal prior. Given that the measurements in this causal scenario stem from different sources, the product is integrated over all possible, remapped visual and auditory stimulus locations, 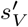 and 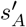:

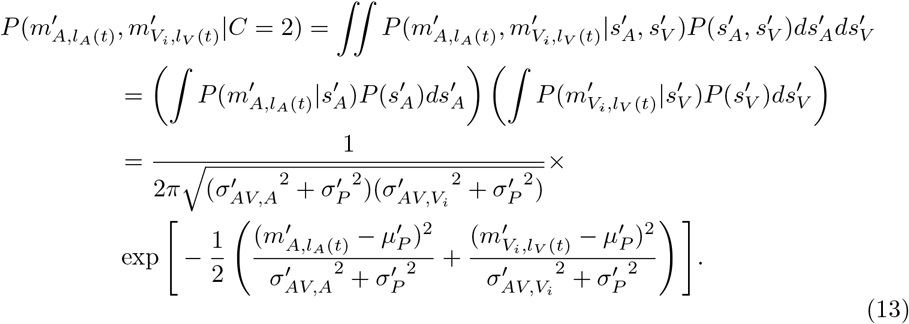

The updates of the shifts are scaled by two modality-specific learning rates, *α*_*A*_ and *α*_*V*_ [62]:

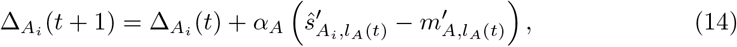

and

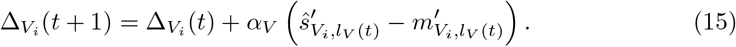

We also tested a version of the model with one supra-modal learning rate (*α* = *α*_*A*_ = *α*_*V*_).

### Formalization of the tasks

As outlined in the beginning, we assumed that visual and auditory stimuli are remapped into a common internal reference frame to which we have no access and this remapping involves modality-specific shifts 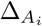 and 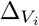. Updates to these shifts were probed in our study by introducing an artificial location-independent spatial discrepancy between visual and auditory measurements. In addition, we assumed that the transformation into an internal reference frame introduces location-specific biases. More specifically, the means of the measurement distributions, the remapped stimulus location in internal space were linear functions of the physical stimulus locations *s*_*A*_ and *s*_*V*_, that is, 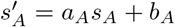 and 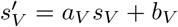 by default and 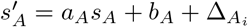 (121) and 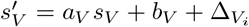 (121) at the end of the recalibration phase. The modality-specific shifts 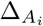 and 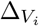 are updated in every trial of the recalibration phase. The updates are independent and thus the location-independent shifts at the beginning of the post-recalibration phase can be written as the sum of the initial shifts and the shift-updates over 120 trials.

The remapping process could bias both measurement functions or just one, but there is no way to empirically distinguish between these possibilities given that we can only measure relative biases. As a consequence and without loss of generality, we set 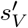 to be equal to *s*_*V*_ before the recalibration (i.e., *a*_*V*_ = 1 and *b*_*V*_ = 0).

### Unimodal spatial-discrimination task

The unimodal spatial-discrimination task was conducted to estimate visual and auditory stimulus reliabilities under unimodal presentation conditions. The task was repeated four times for each participant, once to measure the reliability of the auditory stimulus 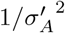 and once to measure the reliability of each of the three types of visual stimuli 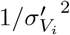.

In every trial, two stimuli of the same type were presented. We begin by describing the auditory version. The standard stimulus was presented straight ahead, at location *s*_*A*,0_, and the test stimulus was presented at one of *N*_*A*_ locations, *s*_*A,n*_, determined by an adaptive procedure. For each pair, the probability, *p*_*A,n*_, of estimating the test stimulus to be located to the right of the standard stimulus is a function of the physical distance between the two stimuli.

We assume that the observer makes the decision by comparing the internal location estimates 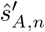 and 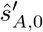,

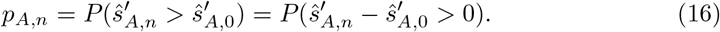

To further specify *p*_*A,n*_, we have to derive the probability distributions of the internal location estimates. Each physical stimulus at location *s*_*A,n*_ results in an internal measurement, 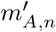. The estimate of the remapped location of this stimulus is the average of the measurement and the mean of the spatial prior, 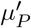, each weighted by their relative reliabilities, respectively:

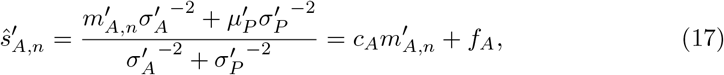

Where

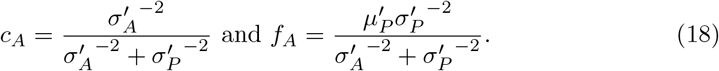

Given that the measurement distribution is Gaussian 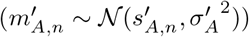 and that the family of Gaussian distributions is closed under linear transformations, the probability distribution of the location estimates of a test stimulus is 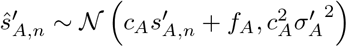 and, analogously, 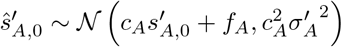 for the location estimates of the standard stimulus. The probability distribution of the difference between the two location estimates 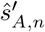 and 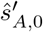 is

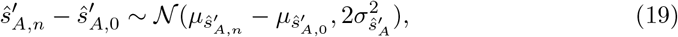

Where 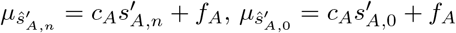, and 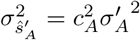. Taken together, the probability of perceiving an auditory test stimulus at location *s*_*A,n*_ to the right of an auditory standard stimulus at location *s*_*A*,0_ is

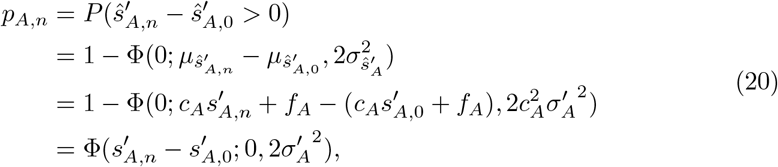

where Φ(*x*; *µ, σ*^2^) is the cumulative Gaussian distribution.

However, as experimenters we only have access to response probabilities as a function of the stimulus locations in physical space. Given that the remapped location of 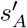 is a function of the physical stimulus location *s*_*A*_, we can rewrite Eq 20 as

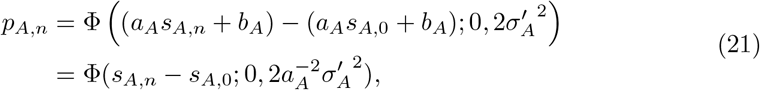

and analogously,

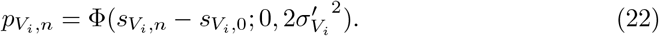

Finally, the model includes occasional response lapses (i.e., random button presses) at rate *λ*, so that the probability of reporting the test stimulus as located farther to the right (*r*_*A,n*_ = 1) was

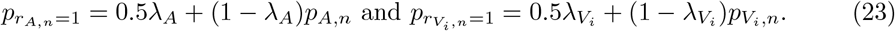

In sum, the probability of reporting the test to the right of the standard stimulus as a function of the relative physical location of the test stimulus is a cumulative Gaussian distribution centered at 0 with variance 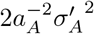 or 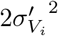. The distribution approaches *λ*_*A*_/2 or 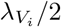 when the test stimulus is far to the left of the standard stimulus, and 1 −*λ*_*A*_/2 or 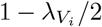 when the test stimulus is far to the right of the standard stimulus (Fig 1C).

### Bimodal spatial-discrimination task

The bimodal spatial-discrimination task was conducted to estimate the relative bias of auditory compared to visual spatial perception introduced by remapping of the measurements into internal space. It was run using the visual stimulus with the highest spatial reliability (*i* = 1). Auditory standard stimuli were presented at four different locations in physical space *s*_*A,l*_, where *l* indexes the auditory location. Guided by a staircase procedure, on each trial *t*, an auditory standard stimulus at location *s*_*A,l*_ was paired with a visual stimulus at one of *N* test locations *s*_*V,n*_, where *n* indexes the finer grid of locations of visual stimuli that were presented during the task. For each pair, the model predicts *p*_*l,n*_, the probability of estimating the visual test stimulus at location *s*_*V,n*_ to be to the right of the auditory standard stimulus at location *s*_*A,l*_.

Analogous to the unimodal spatial-discrimination task, we specify the probability distributions of the internal auditory and visual location estimates as

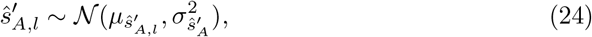

Where

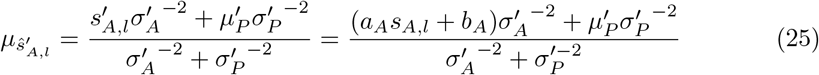

and

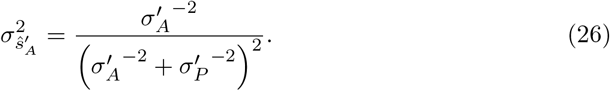

Analogously,

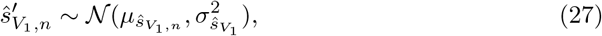

Where

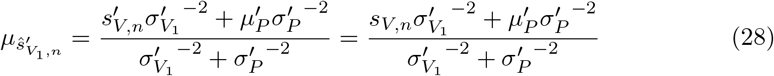

and

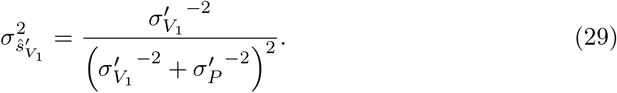

The probability of the observer perceiving a visual test stimulus at physical location *s*_*V,n*_ as located to the right of an auditory standard stimulus at physical location *s*_*A,l*_ is thus:

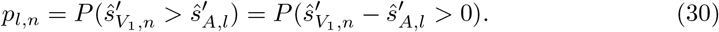

The distribution of this difference is:

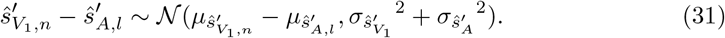

The probability of perceiving the visual test stimulus to the right can then be expressed as

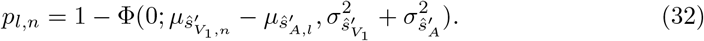

As in the unimodal spatial-discrimination task, the model includes occasional lapses at rate *λ*_*AV*_. Therefore, the probability of reporting a visual stimulus at *s*_*V,n*_ as located farther to the right of an auditory stimulus at location *s*_*A,l*_ (*r*_*l,n*_ = 1) is equal to

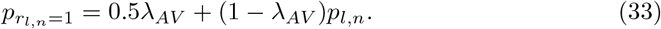

In sum, the probability of reporting the visual stimulus as located to the right of the auditory stimulus as a function of the distance between the two stimulus locations is described by a cumulative Gaussian distribution that approaches λ_*AV*_/2 when the visual test stimulus is located far to the left and 1 − λ_*AV*_/2 when it is far to the right of the auditory standard stimulus.

### The pointing practice task

The pointing practice task was run to familiarize participants with the response device and to provide an estimate of the variability of localization responses in physical space, 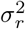, due to sources unrelated to the spatial perception of the stimuli. To achieve this, in this task, participants localized different visual stimuli with minimal spatial uncertainty. Additional noise in the localization responses can, for example, arise from holding the stimulus location in memory before making a response and variability in the use of the response device. Localization of the visual stimuli was tested at eight different locations *s*_*V,o*_, where *o* indexes the stimulus location (*o* ∈ {1, 2, …, 8}). In each trial, participants moved a visual cursor to the perceived location of the stimulus, that is, they confirmed the cursor position as the localization response when the estimated location of the cursor in internal space was identical to the estimated, remapped location of the visual stimulus. The location of the cursor in physical space maps directly to its location in perceptual space, because we assumed the identity mapping for visual stimuli. In addition, we assume that variability in the spatial perception of the highly and constantly visible cursor is negligible. Thus, its internal location estimates should not be biased towards the mean of the spatial prior. In sum, the probability distribution of the confirmed cursor position is centered on the location estimate of the visual stimulus. For the same reasons (identity mapping and negligible spatial uncertainty), in this task, the probability distribution of the stimulus location estimate is centered on the physical stimulus location and response variability should be due only to non-perceptual sources of variability:

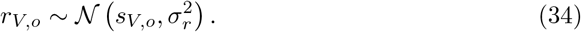

See S1 Appendix: S2 for a model that does not have these assumptions.

### Unimodal localization task

During the recalibration sessions, the unimodal localization task was conducted twice, during the baseline and the post-recalibration phase, to measure the shift in auditory and visual localization responses as a consequence of exposure to spatially discrepant audiovisual stimuli. Stimuli were presented at four locations for each modality, *s*_*A,l*_ and *s*_*V,l*_.

As for the pointing practice task, we assumed that the probability distributions of the localization responses (confirmed cursor positions) in physical space, *r*_*A,l*_ and 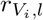, are centered on the location estimates 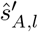 and 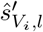 in perceptual space, and that additional unbiased noise corrupts the localization responses. As in the spatial-discrimination tasks (and unlike in the pointing practice task that used the highest-reliability stimulus), the stimulus location estimates are assumed to be biased due to the remapping process and the incorporation of the supra-modal spatial prior. It follows that the probability distributions of the localization responses are

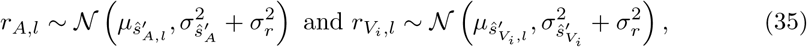

where the terms are defined in Eqs 24–29.

In the post-recalibration phase, the remapping from physical to perceptual space had been updated so that additional shifts 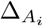 and 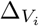, accumulated during the recalibration phase, were incorporated: 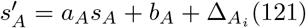 and 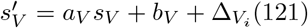. This change in the measurement distributions affects the centers of the location estimates’ probability distributions:

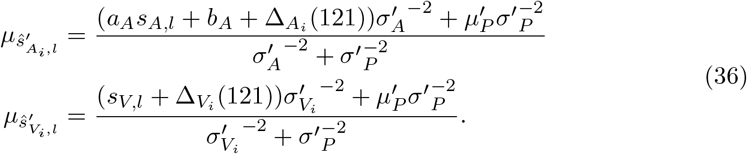

We assumed that the probability distributions of the localization responses are centered on these updated values, that is, we did not implement the updated remapping for the location estimates of the cursor. In sum, localization responses to unimodally presented visual and auditory stimuli have a Gaussian probability distribution that, after the recalibration phase, additionally depends on the final shift updates 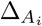 (121) and 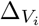 (121).

### Model fitting

For a given model *M*, we fitted the model jointly to the forced-choice responses from the bimodal spatial-discrimination task and to the localization responses from the unimodal spatial-localization task (the baseline and the post-recalibration phases of the recalibration sessions). We fitted the model separately to the forced-choice responses from the unimodal spatial-discrimination task and the localization responses from the pointing practice task to reduce the number of free parameters estimated at once. We did not fit the localization responses from the recalibration phase because the shifts (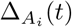 and 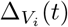) are serially dependent, that is, the size of the shift in trial *t* depends on the size of the shift in trial *t*−1. Given that there is no closed-form solution for the model, we would have needed Monte Carlo simulations to approximate the distribution of location estimates for trial *t* = 1, and for trial *t* = 2, again we would have needed Monte Carlo simulations to approximate the distribution of location estimates given each of these shifts from trial *t* = 1. Because of this serial dependence, the number of necessary samples would have grown exponentially from trial to trial. Thus, it was infeasible to estimate a likelihood of the parameters and the model given the data from the recalibration phase. Instead, we used Monte Carlo simulations to approximate the probability distribution of the final modality-specific shifts 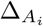 (121) and 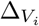 (121) accumulated at the end of the recalibration phase.

All models were fit using a maximum-likelihood procedure. That is, a set of free parameters Θ was chosen to maximize the log likelihood

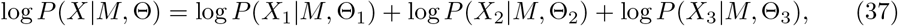

where *X*_1_, *X*_2_, *X*_3_ ⊂ *X* and Θ_1_, Θ_2_, Θ_3_ ⊂ Θ. The first two datasets, *X*_1_ and *X*_2_, and parameter subsets, Θ_1_ and Θ_2_, refer to the unimodal spatial-discrimination task and the pointing practice task, respectively. The third subset comprised data and parameters from three tasks, 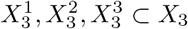 and 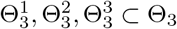, the bimodal spatial-discrimination task, and the unimodal spatial-localization responses from the baseline and post-recalibration phases, respectively. The log-likelihoods of each subset *X*_*i*_ were maximized separately, but the three sets 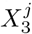 were fit jointly. The three models of cross-modal recalibration only varied with respect to 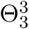.

### Model log-likelihood - unimodal spatial-discrimination task

In the unimodal spatial-discrimination task (auditory session), for each trial *t*, participants were presented with, in random order, a standard stimulus located straight ahead at *s*_*A*,0_ and a test stimulus presented at one of *N*_*A*_ locations *s*_*A,n*(*t*)_. Participants indicated whether the test stimulus was located to the left, *r*_*A,n*(*t*)_ = 0, or to the right of the standard stimulus, *r*_*A,n*(*t*)_ = 1. For each such trial, the likelihood of model parameters given the response *r*_*A,n*(*t*)_ is

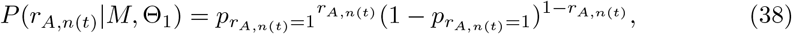

where 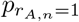 is defined in Eq 23. Thus, the log likelihood given responses across all *T*_1,*A*_ trials is

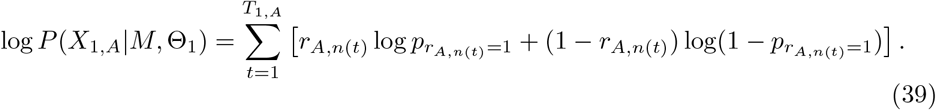

Analogously,

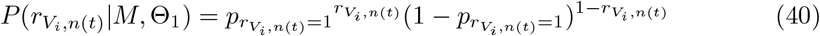

and

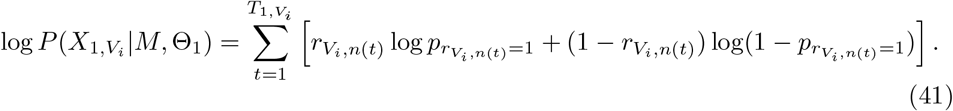

The log likelihood across all four sessions is

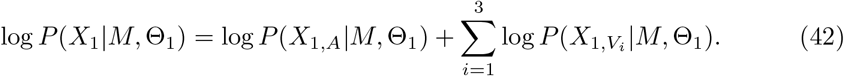

The set of free parameters that were constrained by the binary responses in this task is 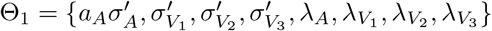.

### Model log-likelihood - pointing practice task

In this unimodal localization task, for each trial *t*, participants were presented with a visual stimulus *s*_*V,o*(*t*)_ with minimal spatial uncertainty. For each such trial, the likelihood of the model parameters given the cursor setting *r*_*V,o*(*t*)_ is 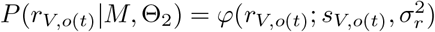 where *φ* refers to the Gaussian probability density. The only free parameter that was constrained by this task is Θ_2_ = {*σ*_*r*_}. The maximum-likelihood estimate of *σ*_*r*_ is

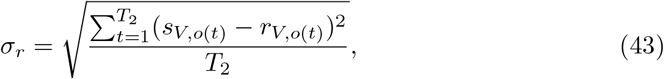

where *T*_2_ and the sum do not include outlier trials.

### Model log-likelihood - bimodal spatial-discrimination task

In the bimodal spatial-discrimination task, for each trial *t*, participants were presented with an auditory and a visual stimulus at locations *s*_*A,l*(*t*)_ and 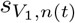, and indicated whether the visual test stimulus was located to the left, *r*_*l*(*t*),*n*(*t*)_ = 0, or to the right, *r*_*l*(*t*),*n*(*t*)_ = 1, of the auditory standard stimulus. For each such trial, the likelihood of model parameters given the response *r*_*l*(*t*),*n*(*t*)_ is

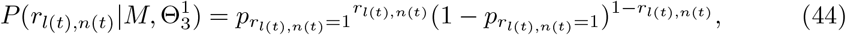

where 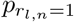 is defined in Eq 33. Thus, the log likelihood given the responses across all 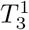 trials is

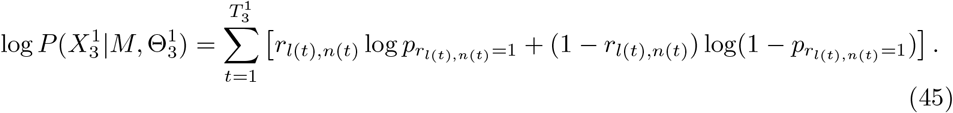

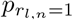 is a function of *p*_*l*(*t*),*n*(*t*)_, which in turn depends on the bias parameters *a*_*A*_ and *b*_*A*_, the parameters of the supra-modal prior over locations 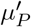 and 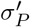, as well as the measurement variances 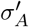 and 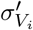 (see Eqs 24–29). Fitting both the bias parameters and the supra-modal prior at once was impossible as they effectively traded off. Thus, we implemented a non-informative supra-modal prior over stimulus locations by setting 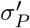 to 100 and 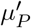 to 0. 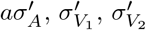 and 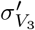 were estimated based on the forced-choice responses from the unimodal spatial-discrimination task. The final set of free parameters that were constrained by the binary responses in this task was 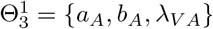 The bias parameters, *a*_*A*_ and *b*_*A*_, were jointly estimated using the data from this task as well as the localization responses from the unimodal spatial-localization task during the baseline and the post-recalibration phases.

### Model log-likelihood - unimodal localization task - baseline phase

In the unimodal localization task, each auditory and each visual localization results in cursor location settings *r*_*A,i,j,l*(*t*)_ and 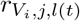 on trial *t* of session (*i, j*) where *i* indicates the visual-reliability condition and *j* the recalibration direction (visual-left-of-auditory or visual-right-of-auditory). The localization responses from this task were modeled as Gaussian-distributed. From these distributions, we can compute the likelihood of a model *M* and the parameter set 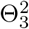 as the Gaussian probability density function in Eq 35 evaluated at the observed localization responses *r*_*A,i,j,l*(*t*)_ and 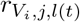 :

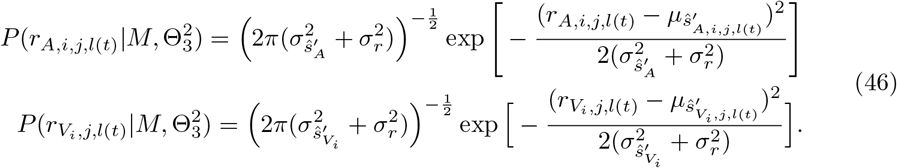

The log likelihood is the sum of the log likelihoods across the trials of all six sessions:

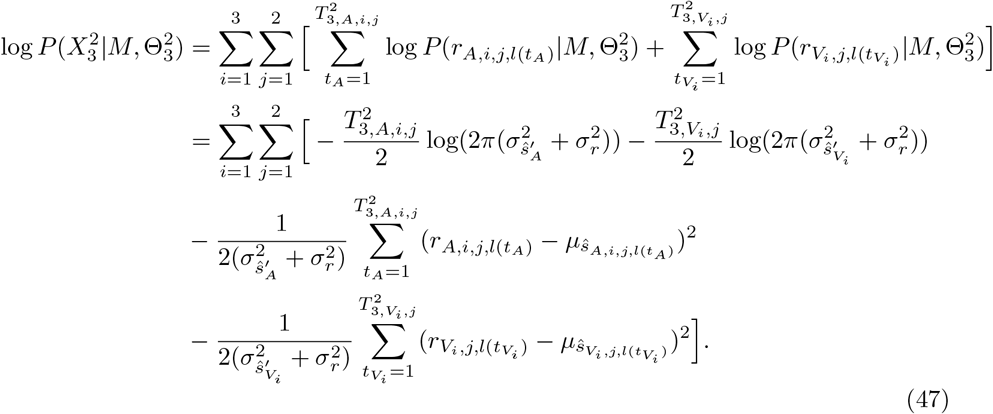

The log-likelihood depends on 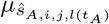 and 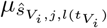, which in turn depend on the bias parameters *a*_*A*_ and *b*_*A*_, the parameters of the supra-modal prior 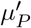 and 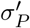, as well as the measurement variances 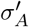 and 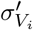 (see Eq 36), and the response noise *σ*_*r*_. Since we chose a flat prior over stimulus locations, the measurement variances were estimated based on the unimodal spatial-discrimination task, and *σ*_*r*_ was estimated based on the pointing practice task, consequently, the actual set of parameters constrained by localization responses from the baseline phase was 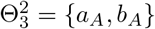. Here, the value of the 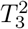 variables and the sums do not include outlier trials.

### Model Log-Likelihood - Unimodal Localization Task - Post-Recalibration Phase

Compared to the baseline phase, localization responses in the post-recalibration phase additionally depend on the updates for the visual and auditory shifts at the end of the recalibration phase, 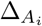 (121) and 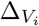 (121) (Eq 36). Since these accumulated shift updates are not accessible to the experimenter, we marginalized over these shift updates to calculate the log-likelihood. For each of the six experimental sessions (*i, j*), the log likelihood of a model *M* and its parameter set 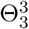 is the integral over 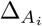 and 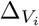 of the likelihood of the final shift updates given the observed data 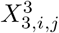, the model *M* and the parameter set 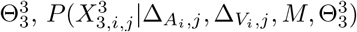, multiplied by the joint probability of the auditory and visual shift updates, 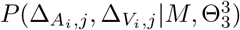 summed across all six sessions

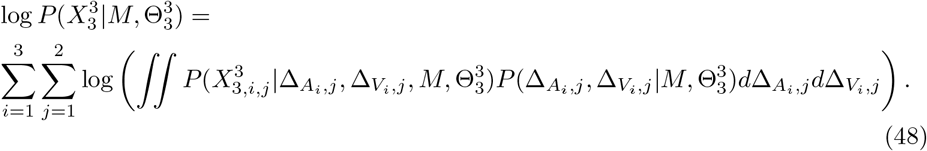

We will describe in the following sections how the joint probability 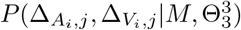 and the log-likelihood log 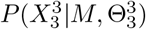 were derived for each of the three models of cross-modal recalibration.

### Log-Likelihood - Reliability-Based Model of Cross-Modal Recalibration

In this model, auditory and visual shift updates have a constant ratio of 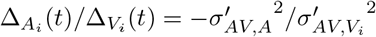 the ratio of the measurement noise variances. Therefore, 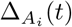 can be rewritten as 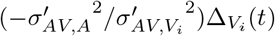, and we can express the likelihood given a single auditory localization response 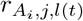 as

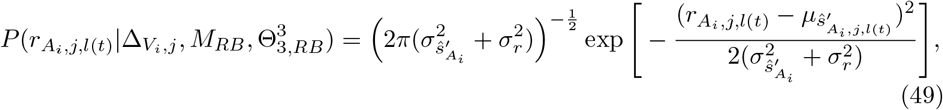

Where

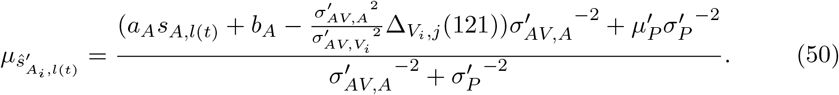

Thus, the joint likelihood 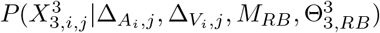 can be written as 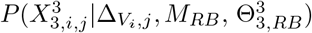. Given that the likelihood depends only on 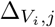 we only need to integrate over 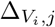 and the log likelihood simplifies to

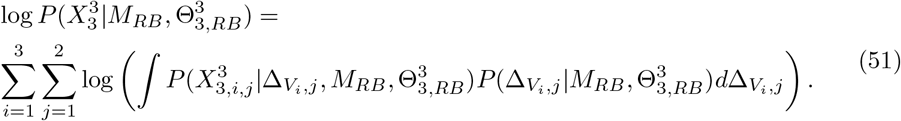

The shift updates 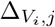 are stochastic because the visual and auditory measurements in each trial of the audiovisual recalibration phase are stochastic. We cannot derive their probability distribution 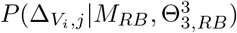 in closed form.

Instead, we used Monte Carlo simulation to approximate this probability distribution. Given the reliability-based model, for each candidate set of parameters 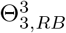, visual-reliability condition *i*, and recalibration direction *j*, we simulated 120 recalibration trials analogous to the audiovisual recalibration phase. We repeated this simulation 1,000 times, resulting in a sample of 1,000 shift updates 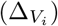 and checked whether the distribution of the 1,000 samples was well fit by a Gaussian with mean and standard deviation equal to the corresponding empirical parameters of the sampled distribution. To do so, we binned the simulated shift updates into 100 bins of equal size and computed the correlation between the observed and predicted number of samples per bin. The resulting value of *R*^2^ was greater than 0.925 in all cases (S1 Appendix: S5). The approximated probability distribution of the shift updates is denoted as 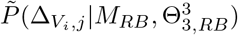.

We approximated the integral in Eq 51 by numerical integration over a region discretized into 100 bins. To ensure that we include enough of the tails of the probability distribution of the shift updates, we set the integration region to be three times larger than the range of the 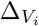 samples, and centered the integration region on that range. Thus, the lower bound, *lb*, is defined as *lb* = Δ_*min*_ − (Δ_*max*_ − Δ_*min*_) and the upper bound is *ub* = Δ_*max*_ + (Δ_*max*_−Δ_*min*_). The numerical integration region was derived separately for each session. The log likelihood is:

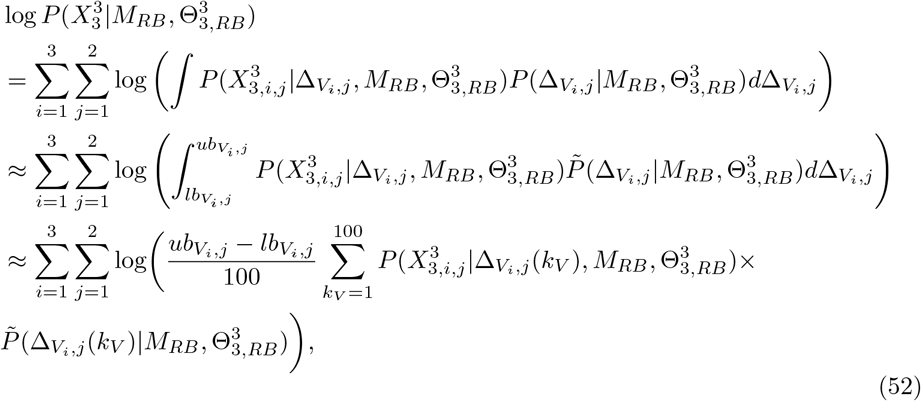

Where

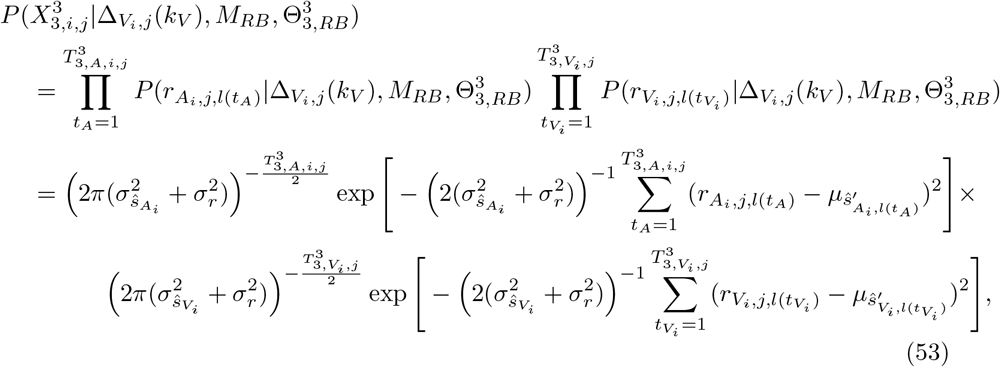

with 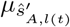 defined in Eq 50 and 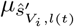 defined in Eq 36. 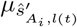 and 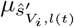 depend on the bias parameters *a*_*A*_ and *b*_*A*_, as well as on 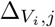, which depends on the measurement variances 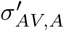 and 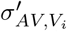 given bimodal presentation and the common learning rate *α*. Note that 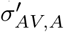 and 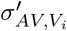 are not directly constrained by data from bimodal trials (because these trials were not included in the model fitting), but estimated based on their influence on the shift-updates. The set of free parameters for this model is 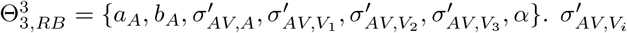 was constrained to be a non-decreasing function of visual-reliability condition *i*, and 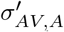 and 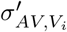 were constrained to be no greater than five times the average values of 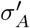 and 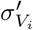 across participants (S1 Appendix: S9). Here, the value of the 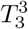 variables and the sums do not include outlier trials.

### Log-Likelihood - Fixed-Ratio Model of Cross-Modal Recalibration

In this model, auditory and visual shift updates have a fixed ratio of 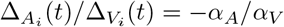 (S1 Appendix: S10). Thus, we can express the likelihood for the fixed-ratio model and parameter set 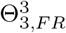 given an auditory localization response 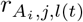 in a similar form to Eq 50:

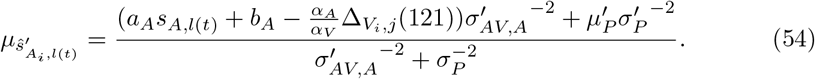

The approximation 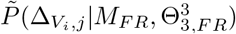 was generated in the same way as for the reliability-based model. The set of free parameters for this model is 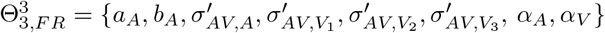. As in the reliability-based model, 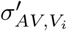 was constrained to be a non-decreasing function of visual-reliability condition *i*, and 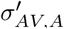 and 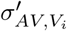 were constrained to be no greater than five times the average values of 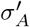 and 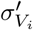 across participants.

### Log-Likelihood - Causal-Inference Model of Cross-Modal Recalibration

For this model, the joint likelihood 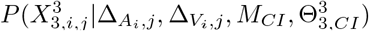 was truly two-dimensional. Thus, we approximated the joint probability of the auditory and visual shift updates, 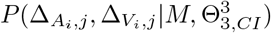 by drawing 1000 samples of shift-update pairs and compared the set of sample pairs to a 2-d Gaussian with the sample mean and covariance as parameters. We again tested whether the two-dimensional Gaussian distribution provided a good fit to the simulated density (defined as *R*^2^ *>* 0.925). If the Gaussian fit was poor, we used a kernel density estimate (Gaussian kernel smoother with *σ* chosen automatically) of the distribution based on the 2-d density of the samples [78, 79]. Overall, the simulated auditory and visual shift updates were very well fit by a bivariate Gaussian, and we rarely used a kernel density estimate (S1 Appendix: S5). We additionally used simulations to verify that our estimates of the partial model log-likelihood 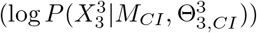 had reasonably small bias (S1 Appendix: S5).

For the causal-inference model, we approximate the log likelihood by numerical integration over a 2-dimensional region of Δ_*A*_, Δ_*V*_ space discretized into 100×100 bins. The upper and lower bounds were determined for both dimensions in the same way as before. The log likelihood is:

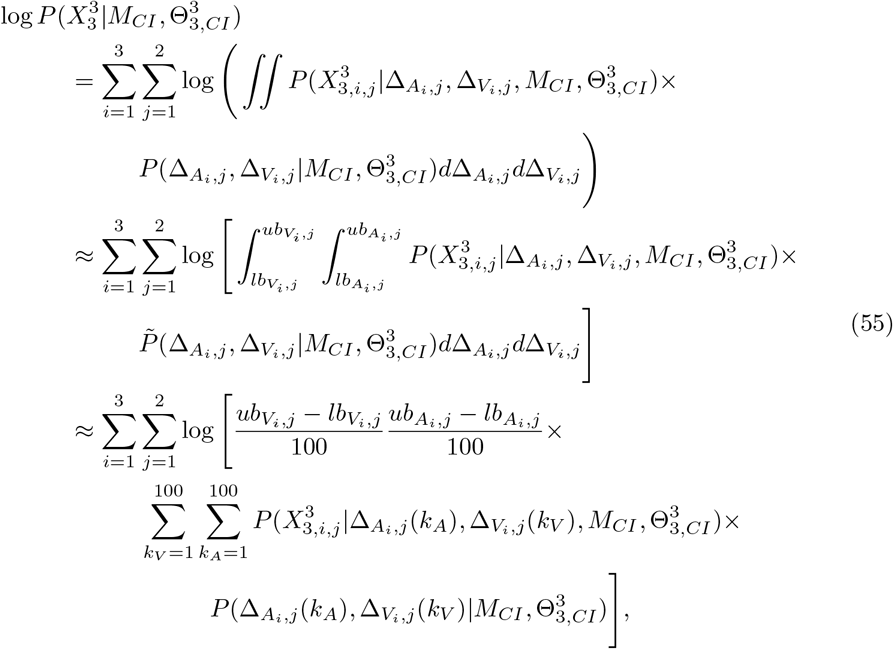

where 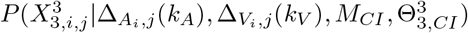 is defined analogously to the reliability-based (see Eq 53) with the exception that 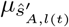 and 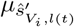 are defined in Eq 36. The set of free parameters used to fit the causal-inference model to the localization responses in the post-recalibration phase is 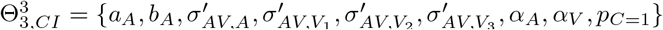 or 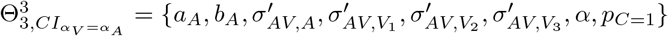.

### Parameter Estimation

For each model, we approximated the set of parameters Θ_1_ and Θ_2_ that maximized the likelihood using the MATLAB function *fmincon* and Python *SciPy*.*optimize* [80], and approximated Θ_3_ using the BADS toolbox [81]. To deal with the possibility that the returned parameter values might correspond to a local minimum, we ran BADS multiple times with different starting points, randomly chosen from a *D*-dimensional grid, where *D* is the number of free parameters in Θ_3_ (see **Table 1** for a summary of the free parameters for each model) and with three evenly spaced values chosen for each dimension. The final parameter estimates were those with the maximum likelihood across all runs of the fitting procedure.

**Table 1.**
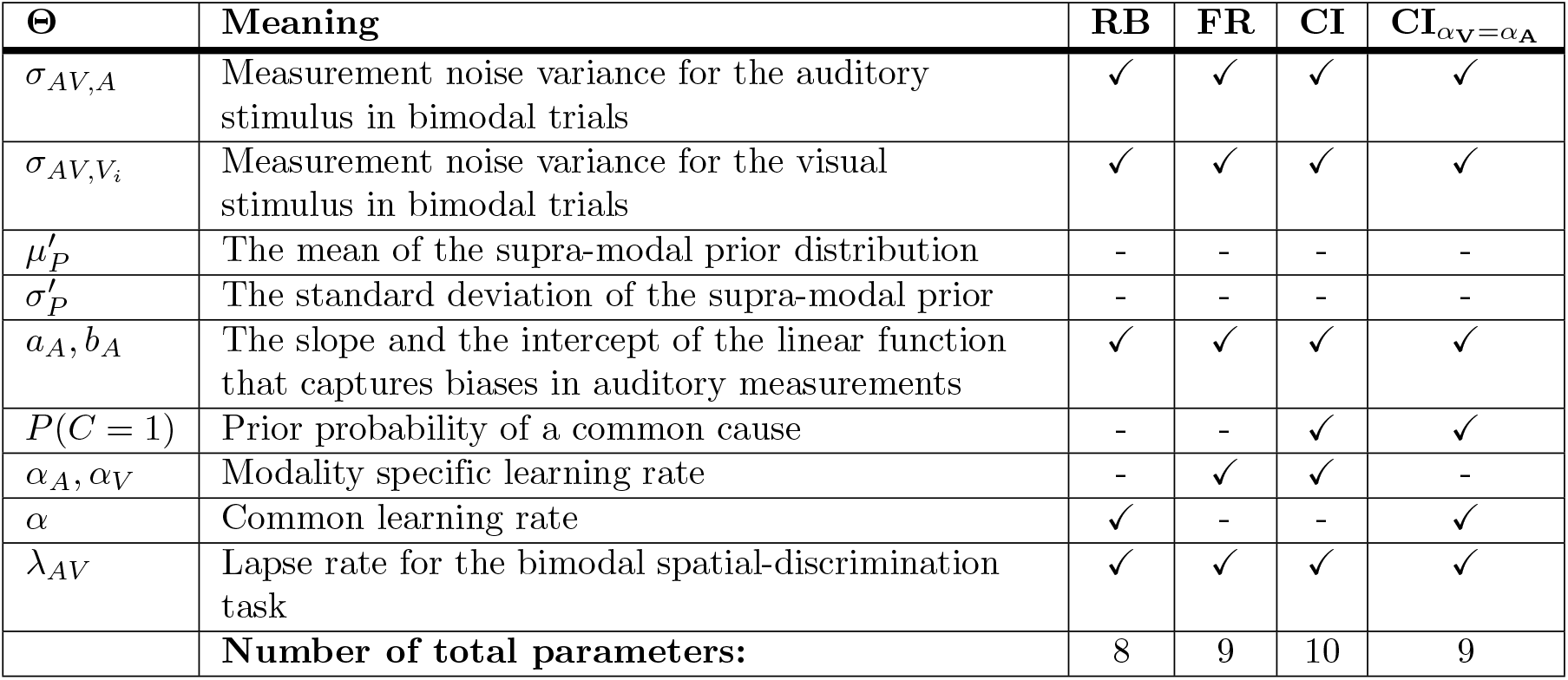
Summary of model parameters.

### Model Comparison

To compare model performance quantitatively, we computed the Akaike Information Criterion (AIC) for all four models [67] and calculated relative model-comparison scores, Δ_AIC_, which relate the AIC value of the best-fit model to that of each of the other models (a higher Δ_AIC_ value indicates stronger evidence for the best-fit model). Models with 0 *<* Δ_AIC_ *<* 2 are weakly supported; models with 4 *<* Δ_AIC_ *<* 7 have considerably less support; models with Δ_AIC_ *>* 10 have essentially no support [82].

## Acknowledgements

We would like to thank Luigi Acerbi for advice on model fitting, and Shannon Locke, Elyse Norton, Hörmet Yiltiz, Elon Gaffin-Cahn, Charlie Burlingham and Antonio Fernandez for support and comments. This work utilized the NYU IT High-Performance Computing resources and services. This work was supported by NIH grant EY08266.

## Author Contributions

**Conceptualization**: Fangfang Hong, Stephanie Badde, Michael S. Landy.

**Data collection**: Fangfang Hong.

**Formal analysis**: Fangfang Hong.

**Funding acquisition**: Michael S. Landy.

**Investigation**: Fangfang Hong.

**Methodology**: Fangfang Hong, Stephanie Badde, Michael S. Landy.

**Supervision**: Stephanie Badde, Michael S. Landy.

**Validation**: Fangfang Hong, Stephanie Badde, Michael S. Landy.

**Visualization**: Fangfang Hong.

**Writing - original draft**: Fangfang Hong, Stephanie Badde, Michael S. Landy.

**Writing - review & editing**: Fangfang Hong, Stephanie Badde, Michael S. Landy.

## Supporting information

**S1 Appendix. Appendix**. Supplemental control experiments, analyses and figures.

